# Decades of native bee biodiversity surveys at Pinnacles National Park highlight the importance of monitoring natural areas over time

**DOI:** 10.1101/462986

**Authors:** Joan M. Meiners, Terry L. Griswold, Olivia Messinger Carril

## Abstract

Thousands of species of bees are in global decline, yet research addressing the ecology and status of these wild pollinators lags far behind work being done to address similar impacts on the managed honey bee. This knowledge gap is especially glaring in natural areas, despite knowledge that protected habitats harbor and export diverse bee communities into nearby croplands where their pollination services have been valued at over $3 billion per year. Surrounded by ranches and farmlands, Pinnacles National Park in the Inner South Coast Range of California contains intact Mediterranean chaparral shrubland. This habitat type is among the most valuable for bee biodiversity worldwide, as well as one of the most vulnerable to agricultural conversion, urbanization and climate change. Pinnacles National Park is also one of a very few locations where extensive native bee inventory efforts have been repeated over time. This park thus presents a valuable and rare opportunity to monitor long-term trends and baseline variability of native bees in natural habitats. Fifteen years after a species inventory marked Pinnacles as a biodiversity hotspot for native bees, we resurveyed these native bee communities over two flowering seasons using a systematic, plot-based design. Combining results, we report a total of 450 bee species within this 109km^2^ natural area of California, including 48 new species records as of 2012 and 95 species not seen since 1999. As far as we are aware, this species richness marks Pinnacles National Park as one of the most densely diverse places known for native bees. We explore patterns of bee diversity across this protected landscape, compare results to other surveyed natural areas, and highlight the need for additional repeated inventories in protected areas over time amid widespread concerns of bee declines.

## Introduction

The importance of bees as critical ecosystem service providers can scarcely be exaggerated. Twenty thousand species of bees worldwide provide the pollination services required for reproduction in 85% of wild and cultivated plants [1,2]. In the United States, the economic importance of bees to agriculture has been valued at up to $14.6 billion annually [3], with $3.08 billion and up to 30% of the U.S. diet specifically credited to the four thousand North American species of native, non-honey bees [4]. Diverse assemblages of native bees have been found capable of enhancing fruit set and yield in the presence of imported honey bees, and of providing adequate pollination for a majority of crops in their absence [5–7]. In natural areas, without the manpower of imported, managed honey bee hives, native bees play a key role in maintaining plant communities that provide soil structure, shelter other invertebrate ecosystem service providers, and establish the base of the food chain [8,9].

Although native bees are often observed pollinating agricultural fields, they seldom nest there. Instead, they rely on nearby remnant patches of semi-natural habitat, a resource that is rapidly disappearing with increasing agricultural intensification, habitat fragmentation, and urban development [10–12]. Despite recognition of natural areas as valuable reservoirs of pollinators [13,14], research on native bee ecology remains concentrated in urban or agricultural settings where baselines may already reflect impacts of degraded ecosystems. Compared to massive honey bee research efforts, progress towards a holistic understanding of how to protect wild bee communities or the habitats they require has not matched their value as pollinators or the known risks they face [15–17].

The relative paucity of research on native bees is due, in part, to the complexity of their biology and behaviors, particularly in wild landscapes. Efforts to monitor wild bees must contend with the ‘taxonomic impediment’ of expertise required to evaluate their vast global biodiversity, and the logistics of sampling a taxon with rapid spatiotemporal turnover, short lifespans, and solitary, elusive habits [18–21]. Unlike many taxa that follow a latitudinal biodiversity gradient [22], bee diversity is highest in xeric and Mediterranean environments, owing to strong seasonal blooms and well-drained soils — features which support a range of foraging specializations and a high temporal turnover of ground-nesting species [19,20,23]. When environmental conditions signal a poor year for host plants, some ground-nesting, specialist bee species can remain underground in diapause for additional years, necessitating multi-year biodiversity monitoring efforts [24]. This fine and irregular partioning of space and time make native bees challenging, time-consuming, and expensive to exhaustively sample in any habitat [25]. Once found, many bee species are difficult to identify even with training and, given reports of functional redundancy within highly-nested pollination networks, the benefit to ecology of doing so may seem unclear [26,27]. However, links between non-random species loss and the stability of ecosystems and mutualistic networks [10,28–34] highlight the merits of species-level bee biodiversity monitoring.

Long-term monitoring of native bee species in natural areas is necessary to reliably assess trajectories of both thriving and struggling native bee communities over time, and to forecast their resilience to future climates and perturbations. Evidence is mounting that climate change affects biotic interactions, increases variability in flowering phenology, and disrupts temporal synchrony between plants and pollinators, potentially impacting plant reproduction and bee access to resources [35–38]. There is a growing need to improve our understanding of the background variability inherent in native bee communities in natural areas in order to contrast that with patterns recorded over time among bee species experiencing a plethora of shifting natural and anthropogenic pressures, including climatic instabilty, shifting habitat phenology, resource depletion, urbanization, and invasion of novel parasites, predators or competitors that may alter ecosystem functioning and the structure of terrestrial communities [36,38,39].

Several large surveys of native bee faunas, particularly in the western United States, have added to current knowledge of the diversity and variability of bee species across space [40–47]. A pair of studies comparing bee faunas from several Mediterranean climate zones concluded that the chaparral habitats of California represent one of the highest global biodiversity hotspots for native bees [48,49]. In the late 1990s, Messinger and Griswold [42] found Pinnacles National Monument in California’s Inner South Coast Range to be one of the most diverse areas known for bees, with 393 bee species discovered in what was then a 68km^2^ area. They attributed this remarkable richness, in part, to Pinnacles’ high floral diversity and habitat heterogeneity [42], features which also make it an ideal place to investigate relationships between native bee community dynamics and environmental variables. In 2002, Pinnacles staff conducted a native bee survey of three changing habitats that added species and a time step to the record of bee biodiversity in the monument.

Fifteen years after that initial species inventory effort and a decade after the smaller survey, we returned to Pinnacles, which became a National Park in 2013, to reinventory its native bee biodiversity and establish a more systematic bee monitoring program [50]. Though several other bee biodiversity studies have spanned multiple years, as far as we are aware, Pinnacles is the only natural region with published results from exhaustive and repeated bee surveys over multiple decades, providing much-needed records of native bee biodiversity over longer periods of time. As such, our study may aid efforts to understand and protect native bee biodiversity in natural areas and help determine restoration goals for bee communities in degraded habitats. Here we seek to (a) present patterns of species occurrence and resource use from three decades of bee species inventories at Pinnacles National Park, (b) examine how bee biodiversity density at this park compares to other published large-scale bee inventories across the United States, and (c) use this literature review and comparison to highlight the need for expanded systematic and repeated bee monitoring efforts in order to understand trajectories and variability of diverse native bee communities over time.

## Materials and Methods

### Site description and collecting history

Pinnacles National Park is a smaller national park, approximately 109km^2^, with a highly dynamic topography. The roughly oval-shaped park is bisected by a high rock-ridge spine running north-south that creates a steep elevational gradient and divides the park into a higher, coastal slope to the west and a drier, lower valley on the east. Initial sampling in 1996 by TLG suggested a rich bee fauna, and motivated the initiation of a more systematic effort to inventory the bee species across the then-monument’s 65km^2^ was undertaken the following year by OMC. This first full inventory spanned 1996-1999 and was conducted along the trail network by opportunistically collecting on a 10-14 day schedule using primarily active (handheld aerial nets) but also passive (pan traps or “bee bowls”) methods during the peak flowering season (locally February through May). Efforts across these years varied in terms of collecting days (as few as 5 or as many as 56 per year), months covered, and locations sampled. In 2002, a passive pan trapping study was conducted by a local park biologist in three grassland plots, with traps placed out every two weeks between March and mid-July, weather permitting. The purpose of this study was to examine changes in bee fauna related to native plant restoration efforts.

In 2005, Pinnacles National Monument acquired an additional 15km^2^ of privately-owned land that expanded the park boundary primarily to the east, but also incorporated some relatively inaccessible lands to the north and south. In 2010, TLG initiated a follow-up biodiversity survey of the bees at Pinnacles, including the new lands to the east. In order to better track temporal trajectories in native bee biodiversity and phenology, we adopted a more systematic park-wide sampling protocol and established long-term monitoring plots where timed, regular collecting events using both nets and pan traps were conducted by JMM across the 2011 and 2012 flowering seasons. The following methods and results are focused on this most recent systematic survey, since a summary of the 1996-1999 inventory has previously been published [42].

### Field methods

For the 2011-2012 re-inventory effort, we established ten 1-hectare long-term plots across a diversity of habitat types and reasonably-accesssible areas of the park. We placed three plots on the western side of the rocky spine divide: two in grasslands and one in a Blue Oak woodland. On the larger, lower-elevation eastern side, we set up three plots in alluvial habitats, two in Live Oak woodlands, and one in a Blue Oak woodland. We also established one plot in a Blue Oak woodland along the high rock spine bisecting the park. One-hectare rectangular plots were roughly 200m by 50m, which fit the constraints of the narrow canyon landscapes. In addition to sampling within plots, we visited areas sampled during the original inventory as well as newly-acquired lands to conduct opportunistic aerial net collecting, and we set out pan traps at the same locations that were sampled using pan traps in 2002 (Fig 1). The geographic coordinates of these ten long-term monitoring locations are included in supplementary materials (S1 Table) and shown in the map of our field site (Fig 1).

**Fig 1.**
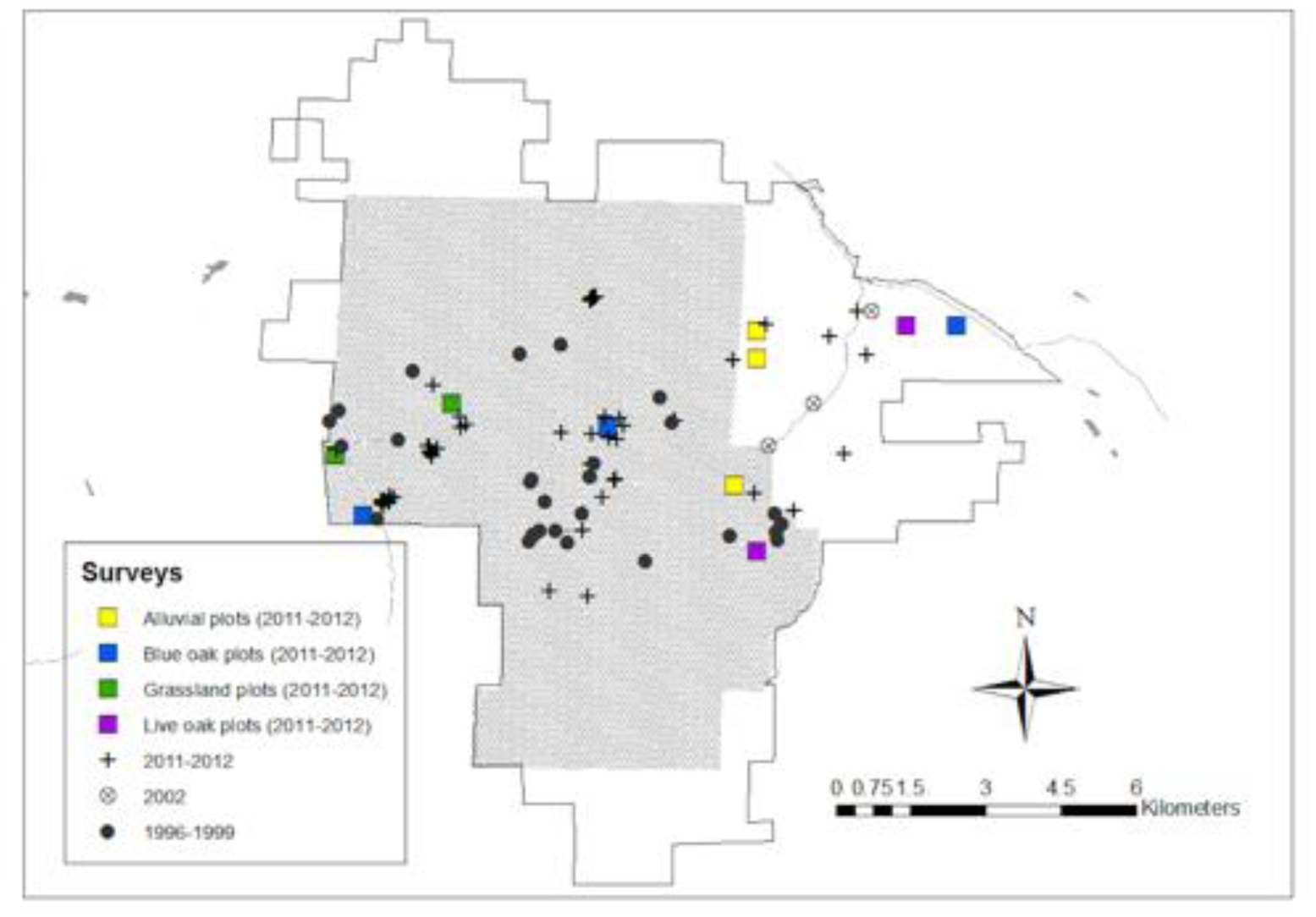
Map of Pinnacles National Park in Monterey and San Benito Counties, California. As a national monument, established in 1908, it grew from 36 km^2^ to 68km^2^, shown by the shaded region. The outlined area encompases lands added in 2005 and represents the current national park boundary (109 km^2^). Locations sampled during the original native bee inventory of 1996-1999 are marked with filled black circles. The three locations where native bees were sampled with pan traps in 2002 are marked by open circles around an ‘x’. For the 2011-2012 survey, plus signs mark sites of opportunistic sampling and colored squares indicate the habitat type and position (not sized to scale) of systematically-sampled hectare plots. Dense chaparral shrubs, steep hillsides, and few trail access points made the northern and southern regions of the park relatively inaccessible for repeated sampling efforts.

Spatially, our collecting extended beyond previous efforts to capture bee biodiversity in three main ways: by traveling off the trail network (along which most collecting was conducted in the 1990s, except for one extensive burned area) for plot and opportunistic sampling, by explicitly establishing repeatedly-sampled plots in a diversity of habitat types across the park, and by venturing into the 15km^2^ of new lands acquired by Pinnacles National Monument in 2005 for both opportunistic and systematic sampling, which had not been done save for one pan-trapping site in 2002 (Fig 1). Temporally, whereas sampling in the 1990s was somewhat irregular, in 2011-12 we sought to capture the full bee community phenology by sampling plots fortnightly throughout the entire flowering season, beginning in February before bee activity began and continuing through late June after most bloom had faded [51].

We sampled all ten plots, typically two per day, every fortnight on days that were mostly sunny, without high winds, and over 15C°. We conducted additional opportunistic net collecting along the trail network or in new off-trail areas in between plot efforts. Immediately before each collecting event, we recorded the ambient temperature, wind speed, humidity, barometric pressure, and a categorical cloud cover value. During plot sampling, two collectors used aerial nets to perform thirty-minute timed collections of all bees visually or auditorily detected in plots at consistent times in both the morning and afternoon. In order to sample the community as evenly and systematically as possible, we walked a steady pace through plots rather than focusing on activity at flowers. We placed all netted bees in vials according to their floral host and collected a voucher plant when the floral host was unknown. At the end of sampling days, we pinned and labeled all specimens and froze them for 48 hours to prevent beetle infestation.

In addition to net collecting, we also set out thirty colored pan traps, a common passive collection method, between 9am and 4pm in each plot on the day we net collected there. Pan traps were made prior to going into the field by spraying 2-oz Solo cups with one of three colors of paint: fluorescent blue, fluorescent yellow, and white, as indicated by the protocol set up for native bee monitoring by Lebuhn et al. [52]. Traps were placed in alternating colors directly on the ground approximately 10m apart in an “X” pattern across rectangular plots and were filled 3/4 full of mildly soapy water to break the surface tension and cause visiting bees to sink to the bottom. At 4pm, we strained insects from the water and immersed them in 75% ethanol until they could be rinsed, pinned and labelled. Data for each pan-trapped specimen includes the color of the bowl from which it was collected.

### Data management and summaries

At the end of the field season, we brought all specimens to the USDA-ARS Pollinating Insect Research Unit (PIRU) in Logan, Utah and incorporated them into its US National Pollinating Insects Collection with the exception of small reference and display collections returned to Pinnacles National Park. Bee identifications were completed by trained experts using Leica dissecting microscopes, taxonomic literature, and the extensive reference collection housed at PIRU (approximately 1.5 million curated bee specimens). After processing all 2011 and 2012 bee specimens, we reviewed all identifications for the Pinnacles bees from the 1996-1999 and 2002 collections (which are also housed at PIRU) to ensure nomenclature was current and consistent with recent inventory identifications. We identified plant vouchers using appropriate keys [53] and guidance from botanists at Pinnacles or the Utah State University Intermountain Herbarium.

We entered field data into PIRU’s existing relational database, assigned corresponding individual ID numbers and barcodes to each specimen, and pinned labels with this information to each bee. We conducted quality checks with multiple people at each step of the curation process. We used SQL and Microsoft Access to query and manage data, and Microsoft Excel, R-Cran statistical package version 0.99.879 or ARC-GIS to clean, arrange, analyze, and map data [54]. Data is either included as supplementary tables or will be deposited with Dryad. Data and code for analysis will be publicly available on Github.

We conducted various summary analyses to asses whether our sampling intensity provided a good characterization of bee biodiversity, and to explore what environmental factors may be related to the bee biodiversity at Pinnacles. We compared species diversity over time by grouping species data across all three sampling collections by year and family and plotting as total values or proportions of total diversity per year. To ascertain whether the recent sampling attempt had captured a sufficient portion of total estimated biodiversity at Pinnacles, we used plot-samplelevel species data to construct a species-accumulation curve with 95% confidence intervals and expected species accumulation values using the ‘vegan’ package in R [55]. We assessed the distribution of bee species data using the Shapiro-Wilk normality test and the relationship between floral richness and bee richness or abundance at the plot-sample level using power-law regression models in the base R package.

### Literature review and study comparisons

To place the bee biodiversity results at Pinnacles National Park in context with those of other bee inventory efforts across the United States, we conducted a literature search for all published studies that reported at least one hundred bee species from natural (non-agricultural, non-urban) areas and methods indicative of an exhaustive, systematic diversity inventory. Using Web of Science and Google Scholar, we identified nineteen published studies that met these criteria, to which we added four unpublished studies that qualify. To allow for a quantitative comparison of relative richness between exhaustive bee surveys, we used a novel metric to calculate biodiversity density along the species-area curve based on the number of species and genera reported in each publication as well as the total size of the area covered, described below. For studies that did not specify the area of land covered, we contacted authors for estimates and/or performed a web search of the study place named to estimate total area surveyed.

Comparisons of the bee species richness over area size reported by different studies was conducted according to Arrhenius’ original description of the species-area relationship as a double logarithmic equation [56,57]:

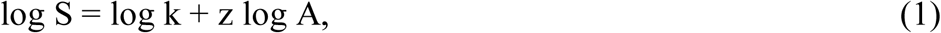

where S represents the number of species recorded in an area of size A, and k and z are constants that may vary with the taxa or habitat assessed.

To quantify the relative richness of studies conducted over different-sized areas and to identify each as recording either above or below the richness per area expected by the relationship defined above, we calculated the distance from each species-area point to the overall log-log regression line calculated according to equation (1) above. We then plotted these observed:expected values in a barplot to compare the relative deviation above or below expected of bee biodiversity values from different studies identified in the literature. These calculations and visualizations were all conducted in R statistical package [54], and data and code are publicly available on GitHub.

## Results

### Pinnacles bee collections over time

Initial trail collecting between 1996-1999 yielded 27,055 bee specimens representing 382 species and 52 genera collected over 125 collector days at 32 different locations within the old monument boundary (Table 1) (differences from results reported by Messinger and Griswold in 2003 are a result of recent taxonomic changes) [42]. The smaller pan trapping study by park biologist Amy Fesnock over 10 days in 2002 yielded 7,255 bees representing 151 species and 38 genera from 3 different locations in the central lowlands of the eastern edge and exterior of the monument boundary. In the recent inventory during the flowering seasons of 2011 and 2012, we captured 52,789 bees over 214 collector days (107 days with two collectors) at 90 different locations across all accessible areas of the park (Fig 1). This effort resulted in a collection of 291 bee species across 45 genera in 2011 and 294 species across 49 genera in 2012 (Table 1a). There was a 79% overlap in species and a 94% overlap in genera between the two years (Table 1b). The preservation and curation of older specimens enabled us to update species determinations from previous inventories based on more recent taxonomic changes to compare and combine biodiversity records across inventory efforts (Table 2).

**Table 1.**
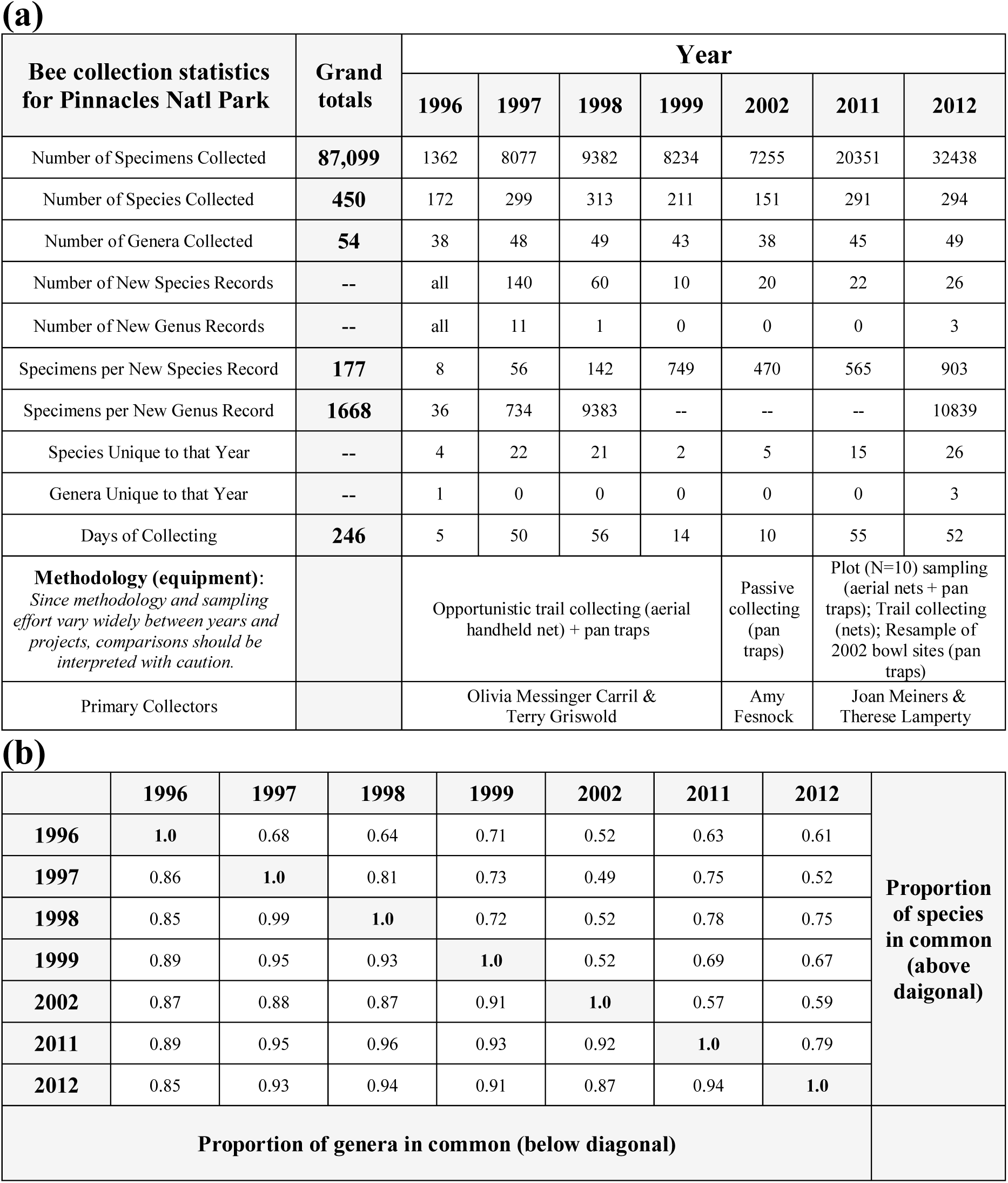
Summary of bee sampling efforts at Pinnacles National Park. (a) Specimen collection statistics by year of sampling. (b) Proportion of overlap between bee species and genera collected during each year of sampling.

**Table 2.** Overview of Pinncles National Park bee biodiversity and comparisons between survey efforts. Numbers of species unique to that survey timeframe are in parentheses. Due to taxonomic changes, updated species determinations, and the addition of data from 2002, some totals differ from those reported in Messinger and Griswold 2003. See S2 Table for additional species details.

| Family | Genus | Number of species in early surveys (1996-1999, 2002) | Number of species in recent survey (2011-2012) | Number of singleton species (represented by only one specimen) | Number of species recorded only in new lands (acquired in 2005) | Cleptoparasitic (C); Oligolectic (O) |
| --- | --- | --- | --- | --- | --- | --- |
| Andrenidae | *Ancylandrena* | 1 (1) |  | 1 |  | O |
|  | *Andrena* | 60 (19) | 49 (8) | 7 | 3 |  |
|  | *Calliopsis* | 8 (2) | 6 |  |  | O |
|  | *Macrotera* | 1 | 1 |  |  | O |
|  | *Panurginus* | 4 (1) | 3 |  |  |  |
|  | *Perdita* | 13 (5) | 10 (2) | 3 | 1 | O |
| Apidae | *Anthophora* | 12 (4) | 8 | 3 |  |  |
|  | *Anthophorula* | 2 | 2 |  |  |  |
|  | *Apis* | 1 | 1 |  |  |  |
|  | *Bombus* | 6 (1) | 5 |  |  |  |
|  | *Brachynomada* |  | 1 (1) |  |  | C |
|  | *Centris* | 1 (1) |  |  |  |  |
|  | *Ceratina* | 11 | 11 |  |  |  |
|  | *Diadasia* | 5 | 9 (3) | 1 |  | O |
|  | *Epeolus* | 3 | 4 (1) |  | 1 | C |
|  | *Eucera* | 9 | 9 |  |  |  |
|  | *Habropoda* | 3 (1) | 2 |  |  |  |
|  | *Melecta* | 3 | 3 |  |  | C |
|  | *Melissodes* | 9 (4) | 8 (3) | 3 | 3 |  |
|  | *Neopasites* |  | 1 (1) | 1 | 1 | C |
|  | *Nomada* | 26 (10) | 21 (5) | 6 | 1 | C |
|  | *Oreopasites* | 2 | 2 |  |  | C |
|  | *Peponapis* |  | 1 (1) |  |  | O |
|  | *Townsendiella* | 2 (1) | 1 | 1 |  | C |
|  | *Triepeolus* | 2 (1) | 7 (5) | 1 | 1 | C |
|  | *Xeromelecta* | 2 (1) | 1 | 1 |  | C |
|  | *Xylocopa* | 1 | 1 |  |  |  |
| Colletidae | *Colletes* | 5 (1) | 5 (1) | 2 |  | O |
|  | *Hylaeus* | 15 (5) | 10 | 2 |  |  |
| Halictidae | *Agapostemon* | 2 | 2 |  |  |  |
|  | *Augochlorella* | 1 | 1 |  |  |  |
|  | *Conanthalictus* | 2 | 2 |  |  | O |
|  | *Dufourea* | 6 | 7 (1) | 1 |  | O |
|  | *Halictus* | 4 (1) | 3 |  |  |  |
|  | *Lasioglossum* | 28 (2) | 29 (3) | 2 | 2 |  |
|  | *Micralictoides* | 2 | 2 |  |  | O |
|  | *Sphecodes* | 10 (5) | 6 (1) | 3 |  | C |
| Megachilidae | *Anthidiellum* | 1 | 1 |  |  |  |
|  | *Anthidium* | 6 (3) | 5 (2) | 1 | 1 |  |
|  | *Ashmeadiella* | 16 (5) | 13 (2) | 2 | 1 |  |
|  | *Atoposmia* | 3 (1) | 3 (1) | 1 |  | O |
|  | *Chelostoma* | 7 | 7 |  |  | O |
|  | *Coelioxys* | 4 (2) | 3 (1) | 1 |  | C |
|  | *Dianthidium* | 5 (1) | 4 | 1 |  |  |
|  | *Dioxys* | 4 (1) | 3 | 1 |  | C |
|  | *Heriades* | 1 (1) |  |  |  |  |
|  | *Hoplitis* | 17 (2) | 15 |  |  | O |
|  | *Megachile* | 18 (5) | 15 (2) | 1 |  |  |
|  | *Osmia* | 38 (6) | 35 (3) | 5 | 2 |  |
|  | *Protosmia* | 1 | 1 |  |  |  |
|  | *Stelis* | 13 (2) | 12 (1) |  | 1 | C |
|  | *Trachusa* | 2 | 2 |  |  |  |
| Melittidae | *Hesperapis* | 2 | 2 |  |  | O |
| **Totals** |  | **417 (95)** | **355 (48)** | **51** | **18** |  |

The combined results from all three inventories document a total of 450 species of bees across 53 genera and all six North American bee families within the modest 109km^2^ of Pinnacles National Park (Table 2). The most recent survey documented 48 new species records for the Pinnacles National Park area and did not recapture 95 species that had been collected in earlier studies (S2 Table). Of the 48 species recorded for the first time in 2011 and 2012, 47 were rare (here defined as represented by fewer than ten specimens), and 20 were singletons (represented by a single specimen) (S2 Table). Thirty of the 48 new species were captured in areas previously sampled, while 18 were only captured in new lands added to the park since previous inventories (Table 2). Overall, 51 of the 450 species were singletons (Table 2), and 95 were present in only one year of sampling, with the majority of these temporally rare species being from the families Apidae and Andrenidae (Fig 2a). The family Megachilidae had the most species present in all seven years of sampling (N=38 out of 68 total) (Fig 2a). Overlap in species lists between years ranged from 49% to 81% and overlap in genera ranged from 85-99% between any two years (Table 1b).

**Fig 2.**
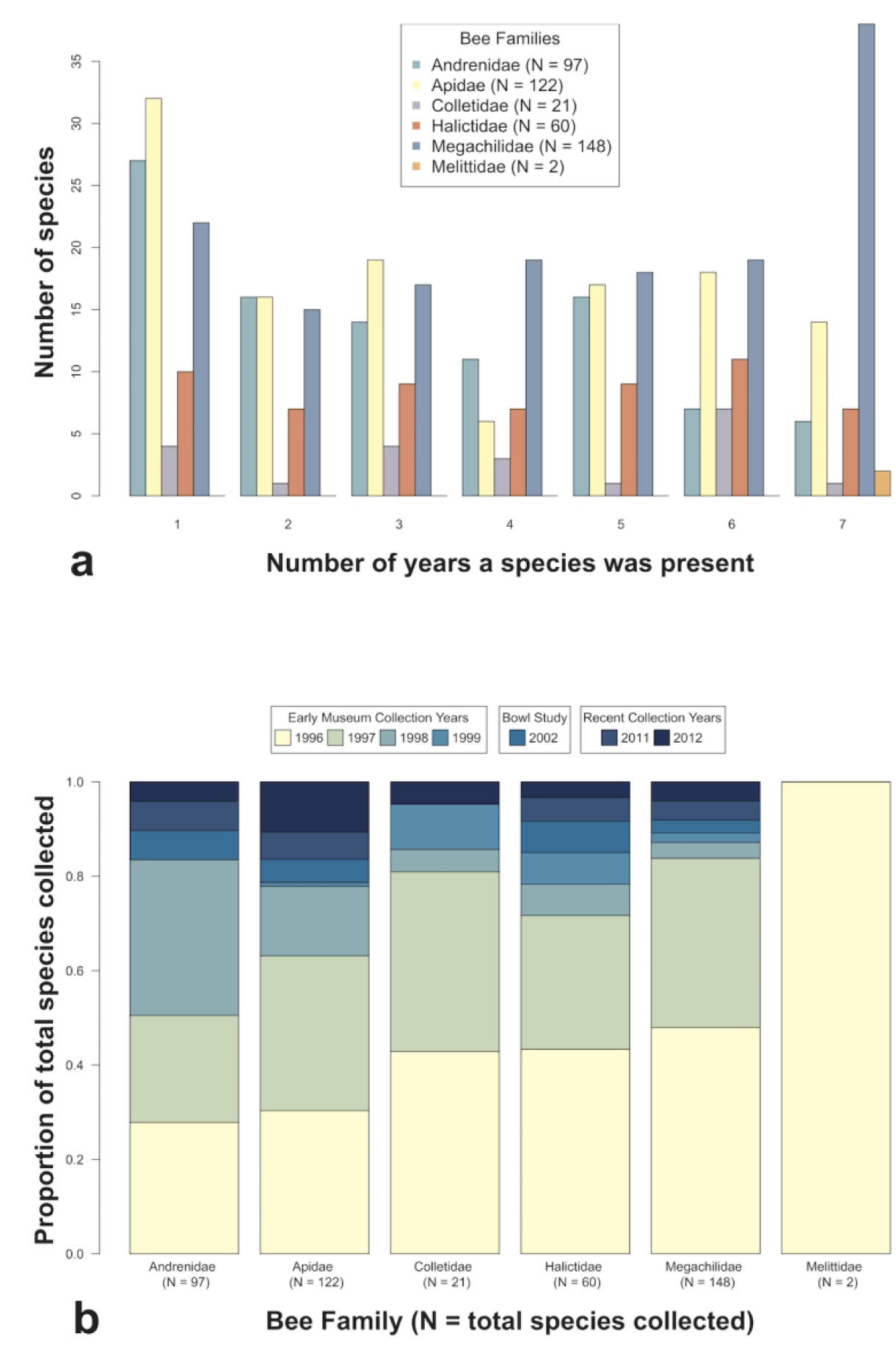
Comparison of bee species collections at Pinnacles National Park over seven years of surveys. (a) Numbers of species in each of six North American bee families represented in up to all seven years of collections. (b) Accumulation over time of number of species collected in each of six North American bee families from each additional year of collecting.

Despite extensive sampling of bee biodiversity within Pinnacles National Monument between 1996-1999, subsequent sampling continued to add species richness to the overall collection (Fig 2b). The 2002 effort added 20 new species to the park list. The 2011 collection netted 22 bee species new to Pinnacles, and the 2012 collection, which sampled mostly the same areas as 2011, resulted in 26 new species and 3 never-before recorded genera within Pinnacles National Park (Table 1a). Between 2 and 26 species were unique to a particular year and not recorded within the park during any of the other six years of surveys. The genus *Ancylandrea* (family Andrenidae) was present only in the 1996 collection and 2012 was the only year that three genera from the family Apidae (*Neopasites, Peponapis*, and *Brachynomada*) were documented (S2 Table). For five out of six bee families, new species were added to the park list nearly every year. Melittidae is represented by only two common species, both of which were collected in the original year of sampling, and in every year thereafter (Fig 2b).

### Recent Pinnacles bee survey details

During the 2011-2012 survey, we completed 150 plot samples across our ten one-hectare plots, eighty in 2011 and seventy in 2012, sampling only on days that were sufficiently sunny, calm, and warm to ensure adequate bee activity for comparisons between plots. In 2011, 80 plot samples conducted over 55 days resulted in between 1 and 2088 bees from an individual plot sample, with a mean of 368 bees per plot per day and a standard deviation of 398. In 2012, 70 plot samples conducted over 52 days resulted in between zero and 1317 bees collected in a day and plot, with a mean of 370 and a standard deviation of 380 bees per plot per day.

A species accumulation curve for the observed rate of capture of the 334 species collected in plots across 150 plot samples shows that our efforts captured a majority of the estimated true bee biodiversity within these areas (Fig 3). The leveling off of the curve at the far right indicates that additional plot sampling would be very slow to yield many more species to the collection, especially for organisms like insects for which observed richness rarely reaches a true asymptote [58]. The prevalence of singleton and doubleton species recorded across many genera illustrates the frequency of rare bee species at Pinnacles National Park, which additional sampling efforts may or may not detect (S2 Table). The blue curve and vertical confidence interval lines indicate the estimated rate of species accumulation for a random community with the same number of species and samples (Fig 3). That the observed curve has an initially steeper slope than expected is indicative of Pinnacles’ rich biodiversity resulting in rapid early accumulation of common species. Expanding collecting efforts into the more remote chaparral habitats in the northern and southern ranges of the park may be more likely to record additional biodiversity without requiring enormous sampling efforts to do so (Fig 1).

**Fig 3.**
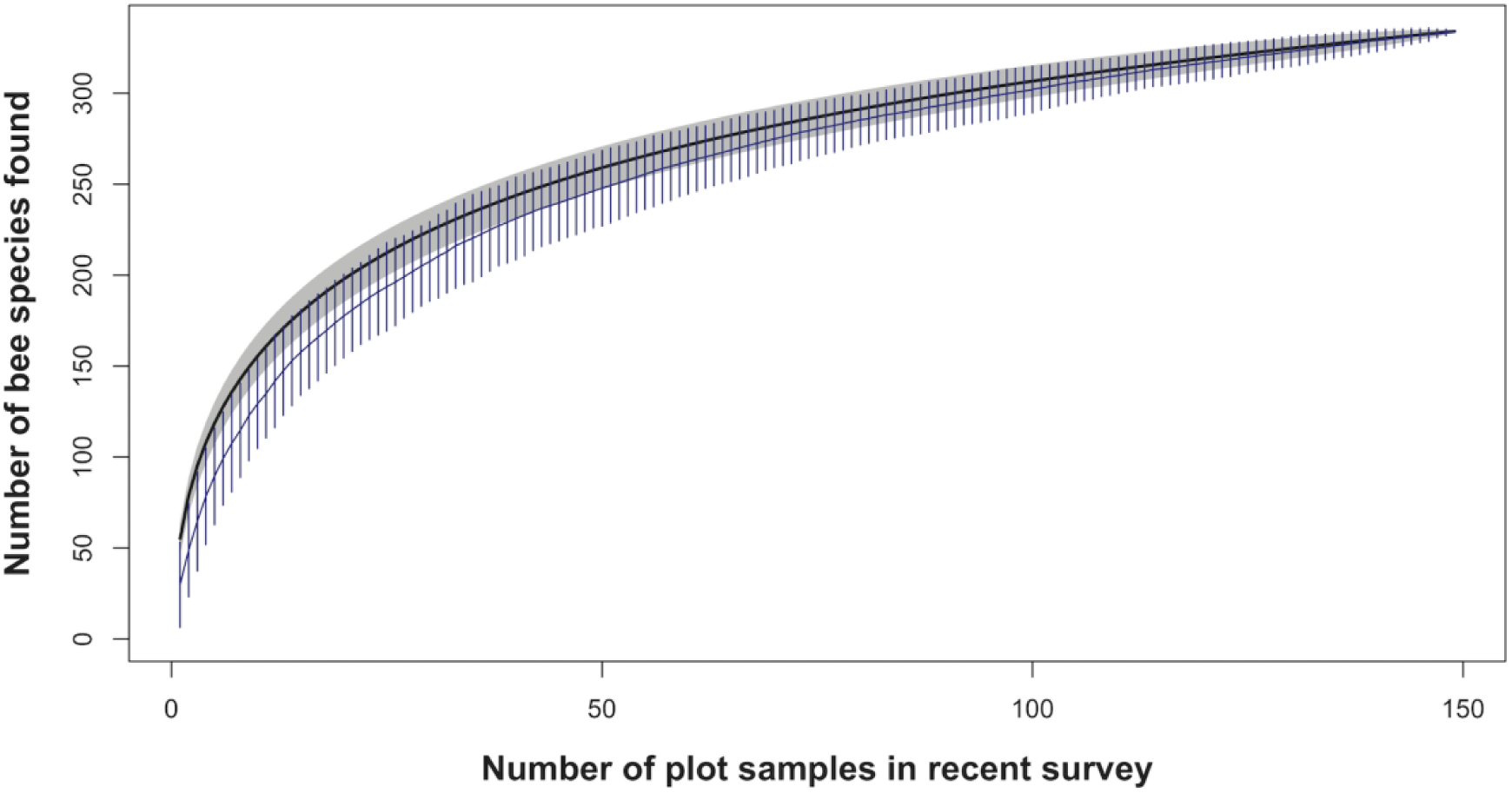
Species accumulation curve. Observed rate of accumulation of 334 species across 150 samples (black line, grey 95% confidence interval bands) compared to an expected rate of species accumulation for a random community with the same number of species and samples (blue line and 95% confidence interval bars).

Bee species richness in 150 plot samples was normally distributed (Shapiro-Wilk normality test, p=0.8) and positively related to the floral richness of bee-visited plants by a power-law linear regression model (Bee Richness = exp(2.79 + 0.38*log(FR)); R^2^=0.37, p<0.01, S1a Fig). To a lesser extent, bee abundance (square-root transformed to normalize distribution) was also significantly positively correlated with the floral diversity of bee-visited plants in plot samples (Bee Abundance = exp(2.26 + 0.23*log(FR)); R^2^ = 0.16, p<0.01, S1b Fig).

Bee abundance, dominance, and floral activity varied between species and the two consecutive years of sampling at Pinnacles National Park. Across all 150 plot samples over two years, *Lasioglossum* (Halictidae) was the most abundant bee genus, followed by *Hesperapis* (Melittidae), *Osmia* (Megachilidae), and *Halictus* (Halictidae). *Oreopasities, Peponapis, Xeromelecta,* and *Townsendiella* (all Apidae) were among the rarest genera collected over the two years of plot sampling; all but *Peponapis* are cleptoparasites.

Between years, rank abundance of the top twenty-five bee species reflects high interannual species turnover, with *Hesperapis regularis* (Melittidae) occupying the top spot in 2011 and only ranking as the fourth most abundant species in 2012 (Table 3a). Similarly, *Osmia nemoris* (Megachilidae) was the most abundant species collected in plot samples at Pinnacles in 2012, after having been ranked fifth most abundant in 2011. Halictidae was the bee family with the highest number of most abundant species in both years, followed by Megachilidae in 2011 and Andrenidae in 2012 (Table 3a).

**Table 3.** Most commonly-collected bees and most bee-popular plants in 2011 and 2012 surveys at Pinnacles National Park. (a) Twenty-five most commonly-collected bee species by rank abundance per year. (b) Twenty-five most commonly recorded plants visited by bees, ranked by popularity with bees per year. See S2 and S3 Tables for the complete taxa lists.

| During the 2011 flowering season |  |  |  | During the 2012 flowering season |  |  |
| --- | --- | --- | --- | --- | --- | --- |
| Bee Family | Genus | Species | Rank Abun. | Bee Family | Genus | Species |
| Melittidae | *Hesperapis* | *regularis* | 1 | Megachilidae | *Osmia* | *nemoris* |
| Halictidae | *Halictus* | *tripartitus* | 2 | Halictidae | *Halictus* | *tripartitus* |
| Halictidae | *Lasioglossum* | *nigrescens* | 3 | Halictidae | *Lasioglossum* | *incompletum* |
| Halictidae | *Lasioglossum* | *brunneiventre* | 4 | Melittidae | *Hesperapis* | *regularis* |
| Megachilidae | *Osmia* | *nemoris* | 5 | Halictidae | *Halictus* | *farinosus* |
| Halictidae | *Lasioglossum* | *incompletum* | 6 | Halictidae | *Lasioglossum* | *nigrescens* |
| Apidae | *Apis* | *mellifera* | 7 | Apidae | *Melissodes* | *stearnsi* |
| Halictidae | *Lasioglossum* | *punctatoventre* | 8 | Apidae | *Apis* | *mellifera* |
| Halictidae | *Halictus* | *farinosus* | 9 | Halictidae | *Lasioglossum* | *brunneiventre* |
| Halictidae | *Lasioglossum* | *sp. 9* | 10 | Halictidae | *Lasioglossum* | *punctatoventre* |
| Halictidae | *Lasioglossum* | *imbrex* | 11 | Apidae | *Eucera* | *actuosa* |
| Apidae | *Ceratina* | *arizonensis* | 12 | Andrenidae | *Panurginus* | *gracilis* |
| Andrenidae | *Andrena* | *aff. cerasifolii* | 13 | Halictidae | *Agapostemon* | *angelicus/texanus* |
| Andrenidae | *Andrena* | *sp.* | 14 | Halictidae | *Lasioglossum* | *sp. 9* |
| Halictidae | *Agapostemon* | *angelicus/texanus* | 15 | Apidae | *Diadasia* | *bituberculata* |
| Andrenidae | *Andrena* | *crudeni* | 16 | Apidae | *Melissodes* | *sp.* |
| Halictidae | *Lasioglossum* | *nevadense* | 17 | Andrenidae | *Perdita* | *distropica* |
| Megachilidae | *Protosmia* | *rubifloris* | 18 | Halictidae | *Lasioglossum* | *sp.* |
| Apidae | *Eucera* | *actuosa* | 19 | Andrenidae | *Andrena* | *aff. cerasifolii* |
| Megachilidae | *Osmia* | *brevis* | 20 | Megachilidae | *Osmia* | *aglaia* |
| Andrenidae | *Panurginus* | *gracilis* | 21 | Megachilidae | *Osmia* | *regulina* |
| Halictidae | *Lasioglossum* | *sisymbrii* | 22 | Halictidae | *Lasioglossum* | *nevadense* |
| Apidae | *Diadasia* | *angusticeps* | 23 | Andrenidae | *Andrena* | *macrocephala* |
| Megachilidae | *Trachusa* | *perdita* | 24 | Andrenidae | *Andrena* | *w-scripta* |
| Megachilidae | *Osmia* | *regulina* | 25 | Apidae | *Ceratina* | *arizonensis* |

(b)
| During the 2011 flowering season |  |  | During the 2012 flowering season |  |
| --- | --- | --- | --- | --- |
| *Plant Name* | *Plant Family* | *Popul. Rank* | *Plant Name* | *Plant Family* |
| *Clarkia unguiculata* | Onagraceae | 1 | *Eriogonum fasciculatum* | Polygonaceae |
| *Adenostoma fasciculatum* | Rosaceae | 2 | *Adenostoma fasciculatum* | Rosaceae |
| *Eschscholzia californica* | Papaveraceae | 3 | *Eschscholzia californica* | Papaveraceae |
| *Clarkia purpurea* | Onagraceae | 4 | *Clarkia unguiculata* | Onagraceae |
| *Chaenactis glabriuscula* | Asteraceae | 5 | *Hirschfeldia incana* | Brassicaceae |
| *Lotus scoparius* var. *scoparius* | Fabaceae | 6 | *Marrubium vulgare* | Lamiaceae |
| *Ranunculus californicus* | Ranunculaceae | 7 | *Eriodictyon tomentosum* | Boraginaceae |
| *Eriogonum fasciculatum* | Polygonaceae | 8 | *Chaenactis glabriuscula* | Asteraceae |
| *Hirschfeldia incana* | Brassicaceae | 9 | *Amsinckia menziesii* | Boraginaceae |
| *Salix exigua* | Salicaceae | 10 | *Salix lasiolepis* | Salicaceae |
| *Lupinus albifrons* | Fabaceae | 11 | *Clarkia purpurea* | Onagraceae |
| *Vicia villosa* | Fabaceae | 12 | *Lasthenia californica* | Asteraceae |
| *Eriodictyon tomentosum* | Boraginaceae | 13 | *Lupinus albifrons* | Fabaceae |
| *Viola pedunculata* | Violaceae | 14 | *Calochortus venustus* | Liliaceae |
| *Quercus agrifolia* var. *agrifolia* | Fagaceae | 15 | *Ceanothus cuneatus* | Rhamnaceae |
| *Lasthenia californica* | Asteraceae | 16 | *Chorizanthe douglasii* | Polygonaceae |
| *Marrubium vulgare* | Lamiaceae | 17 | *Erodium cicutarium* | Geraniaceae |
| *Pholistoma auritum* | Boraginaceae | 18 | *Salix exigua* | Salicaceae |
| *Arctostaphylos pungens* | Ericaceae | 19 | *Penstemon heterophyllus* | Plantaginaceae |
| *Amsinckia menziesii* | Boraginaceae | 20 | *Lotus scoparius* var. *scoparius* | Fabaceae |
| *Ceanothus cuneatus* | Rhamnaceae | 21 | *Baccharis salicifolia* | Asteraceae |
| *Bloomeria crocea* | Liliaceae | 22 | *Vicia villosa* | Fabaceae |
| *Heliotropium curassavicum* | Boraginaceae | 23 | *Malacothamnus aboriginum* | Malvaceae |
| *Erodium brachycarpum* | Geraniaceae | 24 | *Ranunculus californicus* | Ranunculaceae |
| *Salix lasiolepis* | Salicaceae | 25 | *Heliotropium curassavicum* | Boraginaceae |

The most bee-popular plants also varied between years. In 2011, more bees visited *Clarkia unguiculata* (Onagraceae), the host plant of 2011’s most abundant bee, *Hesperapis regularis*, than any other plant (N= 247, compared to 116 bees on this flower in 2012), and *Eriogonum fasciculatum* (Polygoneaceae) was visited by the most bees in 2012 (N = 644, compared to 109 bees on this flower in 2011) (Table 3b). *Adenostoma fasciculatum* (Rosaceae) and *Eschscholzia californica* (Papaveraceae) maintained their positions as the second and third most bee-popular plants, respectively, in both years of collecting. Floral species from the Boraginaceae family dominated the list of top twenty-five most bee-popular plants in 2011 and tied with Asteraceae and Fabaceae for most bee-popular family in 2012 (Table 3b). A broader examination of bee metrics across different habitat types can be found in Meiners 2016 [51].

### Pinnacles bee biodiversity in context

To assess the bee biodiversity density at Pinnacles relative to other locations, we used literature searches and expert opinions to compile a list of 23 studies within the United States that matched our criteria for comparison (N > 100 species, extensive inventory-style sampling in a natural area) (Table 4). It is worth visualizing that, while efforts to survey native bees have increased in recent years, these published inventories still only cover a small proportion of natural areas and habitat types across the United States, and thus offer only a small window into the status of native bees across the country (Fig 4).

**Table 4.** Bee biodiversity density results for all known native bee inventory projects with at least 100 species in natural or semi-natural areas across the United States (N= 23).

| Study location | Study daates | Species | Approx. total area (km<sup>2</sup>) | References |
| --- | --- | --- | --- | --- |
| Grand Staircase Escalante National Monument, UT | 2000-2003 | 656 | 7,610 | [43] |
| Clark County, NV | 1998; 2005, 2006 | 598 | 20,487 | [40] |
| Yosemite National Park, CA | 2006-2009 | 554 | 3028 | *pers. comm. T. Griswold* |
| Pinnacles National Park, CA | 1996-1999; 2002; 2011-2012 | 450 | 109 | *present results & [42]* |
| San Bernardino, AZ<sup>a</sup> | 2000-2007 | 383 | 1,088<sup>a</sup> | [23] |
| Carlsbad Caverns National Park, NM | 2010-2011 | 364 | 189 | *pers. comm. T. Griswold* |
| Curlew Valley, ID | 1969-1974 | 340 | 4,999 | *[59] & updated totals by pers. comm. T. Griswold* |
| San Rafael Desert, UT | 1979-1992 | 333 | 5,180 | [60] |
| Mojave National Preserve, CA | 1975-1995 | 305 | 6,475 | *pers. comm. T. Griswold* |
| Black Hills of SD and WY | 2010-2011 | 290 | 12,950 | [46] |
| Carlinville, IL<sup>a</sup> | 1884-1916 | 288 | 256<sup>a</sup> | [23] |
| Plummers Island, MD<sup>b</sup> | 1920s-2006 | 232 | 0.15 | [61] |
| MPG Ranch, MT | 2013-2015 | 229 | 39 | [62] |
| Indiana Dunes, IN | 2003, 2004; 2010 | 204 | 60 | [63] |
| Albany County, WY<sup>a</sup> | 1995-1996 | 200 | 11,160<sup>a</sup> | [64] |
| Palouse Prairie, ID | 2012-2013 | 174 | 2,122 | [65] |
| Dugway Proving Ground, UT<sup>a</sup> | 2003, 2005 | 163 | 3,243<sup>a</sup> | [45] |
| Channel Islands, CA | Not specified | 154 | 904 | [66] |
| Black Rock Forest Preserve, NY | 2003 | 144 | 15.5 | [47] |
| Tonasket Ranger District, WA<sup>a</sup> | 2004 | 140 | 1,678<sup>a</sup> | [67] |
| Black Belt Prarie, MS<sup>a</sup> | 1991-2001 | 118 | 803<sup>a</sup> | [68] |
| Archibold Biol. Station, FL |  | 113 | 21 | [69] |
| Hattiesburg, MS<sup>a</sup> | 1943-1944 | 104 | 140<sup>a</sup> | [70] |

<sup>a</sup>Area sizes not specified by publication or through author communications were estimated by calculating known size of map area named in study.
<sup>b</sup>The Plummer's Island study was eliminated as an outlier in the species-area relationship shown in Fig 5 because of its extremely restricted area size sampled compared to other studies.

**Fig 4.**
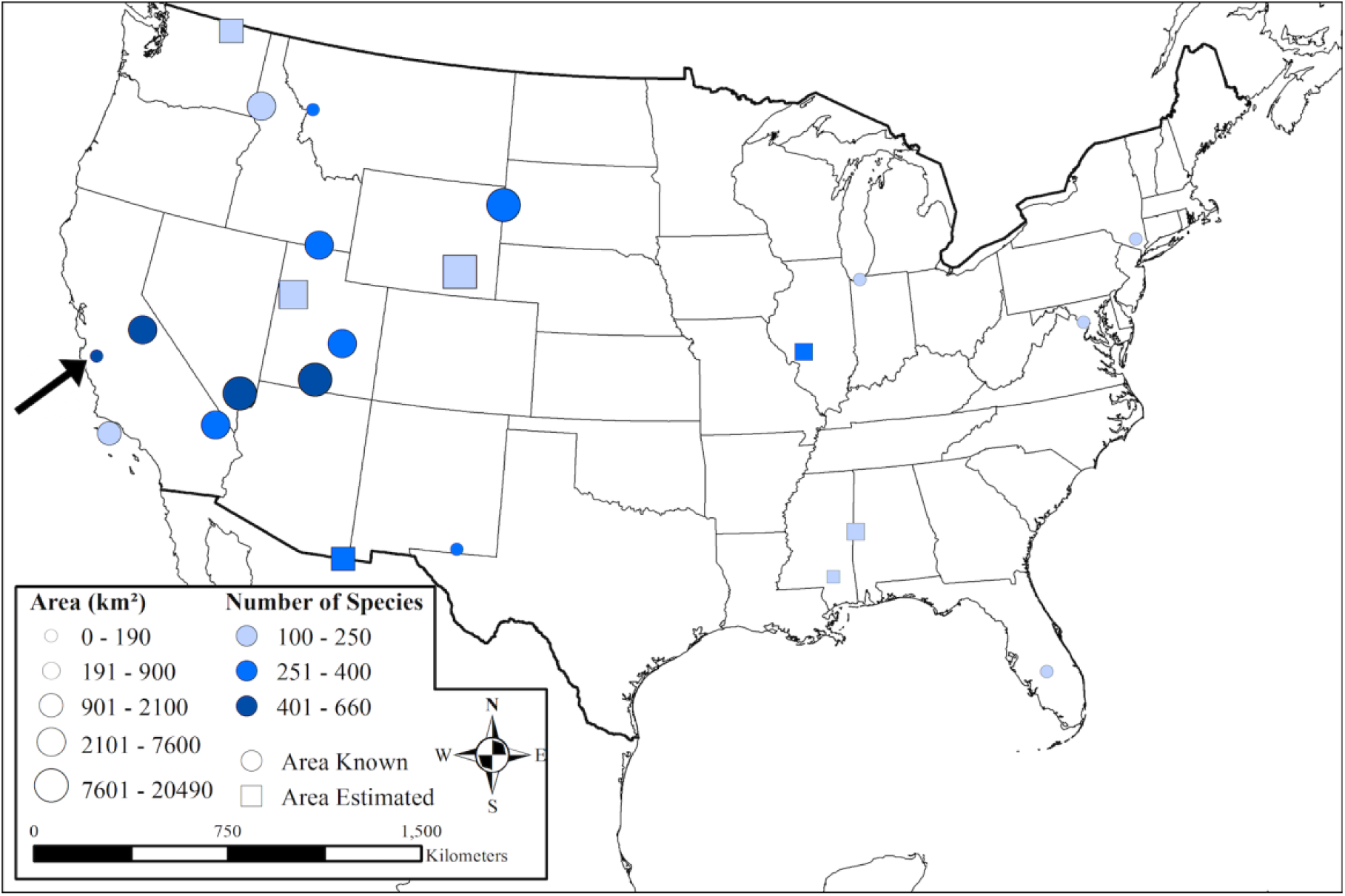
Map of the location, size, and number of bee species recorded for all exhaustive bee inventory efforts undertaken across the United States for which data is published or reported. The black arrow points to Pinnacles National Park. See Table 4 for project details.

Without controlling for the area sampled, Pinnacles’ 450 bee species place it fourth among 23 completed studies reporting high numbers of bee species within a natural area. Studies with more total bee species include Grand Staircase Escalante National Monument, where OMC recorded 656 different species of bees between 2000-2003 [43], a study conducted by TLG in Clark County, Nevada that documented 598 bee species over three years [40], and an unpublished study in Yosemite National Park in the mid-2000s that found 554 species (*Griswold, unpublished data*). A variety of additional systematic inventories conducted in natural lands also report high bee biodiversity, including 393 bee species found over seven years in San Bernardino Valley, Arizona [23], previously thought to have the highest biodiversity of native bees by area.

A meaningful biodiversity comparison between this list of bee inventories is hindered by the vastly different areas each covers. A more direct comparison of the biodiversity of different surveys requires accounting for these differences in area. Because species richness does not scale linearly with spatial area [71,72], we plotted a power-law species-area relationship based on the reported species richness and area covered by known bee inventories (Table 4) to calculate which of the 23 listed studies found lower-than-expected bee richness based on their size and which studies were likely true hotspots of native bee biodiversity (Fig 5).

**Fig 5.**
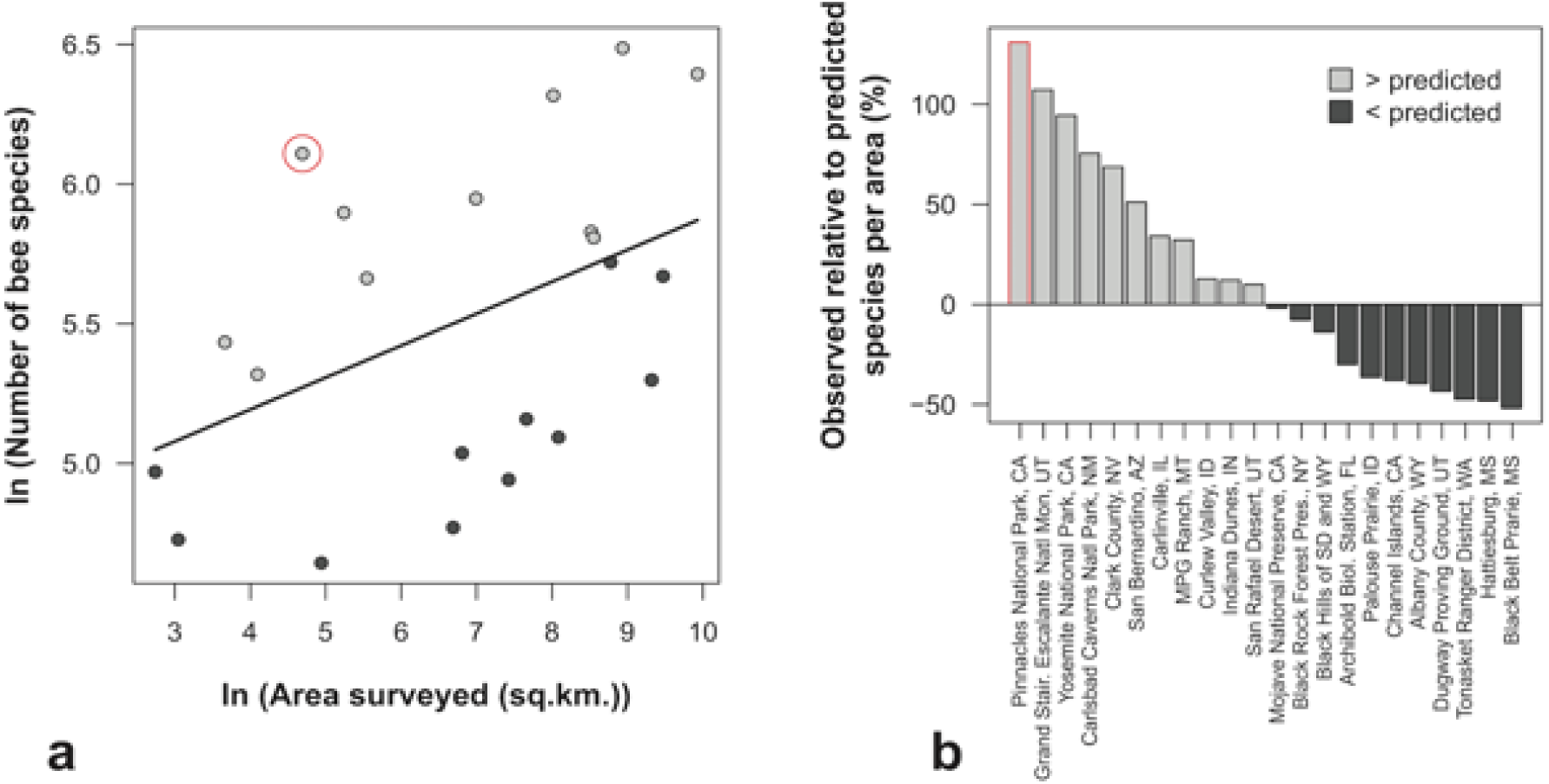
Species-Area relationships and trend line for all major, exhaustive bee inventory studies conducted in the United States in natural or semi-natural habitats. (a) The black trend line delineates expectations for how the number of species will increase with increasing area size based on the (log-transformed) species-area relationship. Studies above the trend line (grey points) recorded more bee species than expected for the area of the site; those below the line (black points) recorded fewer bee species than might be expected on average for that size area. Pinnacles National Park is circled in red. (b) Barplot of the difference in the number of bee species observed in each study relative to the number of bee species predicted by the trend line plotted in panel (a). Pinnacles National Park is outlined in red. Study details are listed in Table 4.

Based on this difference between observed and expected species richness per area (the positive or negative distance of the point to the trend line in Fig 5), we conclude that Pinnacles National Park is home to the highest bee biodiversity per area surveyed of any published or known exhaustive bee biodiversity survey (with over 100 species) in natural areas across the United States. Grand Staircase Escalante National Monument (GSENM) also contains more bee biodiversity than would be expected by even its vast size, as does Yosemite National Park; Carlsbad Caverns National Park; Clark County, Nevada; San Bernardino, Arizona; Carlinville, Illinois; MPG Ranch, Montana; Curlew Valley, Idaho; Indiana Dunes, Indiana; and San Rafael Desert, Utah. Studies that reported bee biodiversity lower than what would be expected by our species-area relationship included Black Belt Prairie, Missouri; Hattiesville, Missouri; Tonasket Ranger District, Washington; and the Black Hills of South Dakota and Wyoming, among other natural areas (Fig 5, Table 4). Many more studies will be necessary to fill in the map of bee biodiversity in natural areas (Fig 4) and interpret how the bee species-area relationship relates to ecosystem, climate, or habitat stage (Fig 5).

## Discussion

Wild, native bees are key ecosystem service providers in both natural and agricultural landscapes [5–7,73]. Compared to the unstable European honey bee, on which United States agriculture is heavily dependent, little is known about the four thousand North American species of native bees, who may also be vulnerable to the same parasites, pesticides, and habitat modification plaguing the honey bee [3,16,17,34,74,75]. One of the reasons for this lack of attention to native pollinators is the expense, time, and skill required to collect and identify native bees, which are spatiotemporally variabile, short-lived, diverse in their taxonomy and nesting habits, and often difficult to see. Even when extensive bee inventories are conducted at intensities and intervals sufficient to capture local diversity in native bees, our literature review found that they are rarely replicated later, resulting in few datasets that allow for robust assessment of trends in native bee populations over ecologically relevant time scales.

With three separate inventories conducted over three decades, the native bee inventory efforts at Pinnacles National Park in the Inner South Coast Range of California represent an exception to this lack of temporal knowledge. Combined results from seven years of sampling suggest that Pinnacles National Park may harbor the highest density of bee species currently known anywhere in the United States, and potentially the world, since California is already recognized as a global bee biodiversity hotspot [20]. In comparison to Pinnacles’ 450 species across an area of 109km^2^, only 388 species of bees have been recorded in the state of Wisconsin and only 40 species on the entire two large islands of New Zealand [76,77]. The closest comparison by habitat type outside of the United States may be a survey conducted 1983-1987 over a Mediterranean area of unspecified size outside Athens, Greece that reported 661 species of bees [78]. A survey of seven California urban areas recorded between 60 and 80 total bee species [73]. However, the fact that substantial species diversity was added to the bee inventory list for Pinnacles even after five prior years of surveys (Figs 2b and 3) suggests that inventories in other locations over shorter timespans may grossly undercount rare species.

Our comparison of the bee biodiversity at Pinnacles with other exhaustive bee surveys conducted in the continental United States supports previous assertions that Pinnacles National Park is home to an expectionally high density of bee species. We attribute the extraordinarily rich bee fauna of Pinnacles National Park to its Mediterranean climate, steep environmental gradients, and high habitat heterogeneity, the last of which has been found in other research to be a stronger predictor of species richness than the species-area relationship [79,80]. Habitat heterogeneity can occur over both space and time. Mediterranean habitats, including those at Pinnacles, are known for rich ‘flash-bloom’ cycles during spring months, followed by hot, dry summers and mild, wet winters, an environment that tends to support a high biodiversity of many taxa by creating many temporal habitat niches [9,81]. Among bees, the rapid turnover of floral resources in these areas may favor solitary species, whose shorter flight periods and more specialized foraging behaviors may allow many species to coexist in a single area, as each occupies a narrower temporal and foraging niche space than longer-lived social or generalist species, which are more common in temperate areas [19,23]. This variability in bee species over time at Pinnacles (Fig 2a) underscores the the importance of long-term sampling to meet the research challenge of detecting the signal amidst the noise of bee community variability [82].

Across space, habitats at Pinnacles change rapidly from the western, coastally-influenced slopes, up the 500m elevational gradient to the rock ridge, and down the different aspects and microclimates of the drier east side. Pinnacles spans several fault lines, the geologic movements of which may have contributed to its elevational variation and broader array of soil types than would typically be found in such a small area [83]. Perhaps because of this soil heterogeneity, Pinnacles is also considered to be a transitional zone between the floral ecotones of northern and southern California [84] and boasts a plant list of nearly 700 species, many of them flowering [85]. We found bee richness to be highly correlated with the richness of bee-visited angiosperms on any given day and site at Pinnacles (S1 Fig), which corroborates results from previous studies [9,43]. Indeed, our conclusion is that the extraordinary diversity of native bees at Pinnacles is a function of the dynamic climate, rich wildflower flora, and landscape patchiness creating a wide array of spatiotemporal habitat niches. These factors may allow more diverse bee communities to coexist across space than has been found anywhere else.

The unparallelled biodiversity of native bees at Pinnacles National Park is especially intriguing given its juxtaposition with nearby agricultural intensity. Salinas Valley, at the doorstep of Pinnacles National Park, produces most of the strawberries, tomatoes, spinach, lettuce, celery, and garlic for the country, along with many smaller crops. Many of the lands surrounding the park that are not irrigated for crops are grazed by cows, which may reduce available floral diversity for bees [86]. Native bees are most diverse in natural, undisturbed areas, proximity to which has been linked to crop pollination success because of the constant influx of wild pollinating insect populations into arated lands inhospitible to long-term residence [11,13]. Agricultural habitats fail to support diverse native bees due to impacts of pesticides, nutritional deficits resulting from monocultures offering only one type of bloom, and practices of tilling and turning over the soil where many native bee species overwinter [5,30,87]. The native bees known to pollinate crops persist not within the fields but in nearby patches of natural, uncultivated land. California has increased efforts to restore habitat for wild bees in agricultural lands. But less attention has been paid to bee source populations in adjacent natural areas, even though source-sink dynamics have recently been determined to influence bee population sensitivity to decline [88]. To date, no measures of bee exchange between Pinnacles and nearby croplands are available, but such data would help define the beneficial halo of bee biodiversity hotspots.

If Pinnacles National Park is indeed a biological refuge for native bee populations within a highly-altered landscape, it will be even more important to track trends in its bee biodiversity over time. Our establishment of ten 1-hectare plots and repeatable methodology will facilitate ongoing monitoring activities and better comparisons of bee biodiversity and population stability over time than are currently possible. During 2011 and 2012, we recorded 355 species of bees at Pinnacles National Park, 48 of which were new records for the park. Initial inventories in the 1990s recorded 382 species, 95 of which we did not encounter during the recent inventory. After six prior years of sampling and a clear leveling of the species accumulation curve, we still recorded three new genera in 2012. These results illustrate the difficulty in deciphering ecological trends from inventories conducted using different methods or in different locations. Long-term, systematic monitoring studies in consistent locations will enable improved understanding of species turnover, range extensions (invasions), local extinctions, baseline states, and how to differentiate natural community variability from bee biodiversity decline, a question we consider a research priority towards assessing pollinator trajectories.

The need for multi-year, temporally replicated bee surveys to better quantify trends and declines in native bees over time is further highlighted by the recent increase in the use of chronosequences, which substitute space as a proxy for time in restored habitats to model changes in native bee dynamics [89,90]. This is a clever approach but increasing efforts to repeat surveys using the same methodology in the same natural areas over actual timespans would be better. Spatial coverage of published bee inventory studies is sparse (Fig 5), and temporal coverage is worse. Expanding long-term bee biodiversity monitoring to additional habitats and supporting the museum work and collection maintenance that enable temporal comparisons will bolster our chances of protecting native bees and agricultural stability.

## Conclusions

Here we reported details of the third extensive bee inventory effort at Pinnacles National Park in California over multiple decades in order to share ongoing findings from a native bee biodiversity hotspot and to highlight the need for additional studies that evaluate temporal trends among pollinators. We are the first to compile and compare similar information on native bee biodiversity from published surveys of natural areas across the United States. With 450 species of native bees, we found that Pinnacles houses a higher density of species than any other natural area studied or than would be expected by the species-area curve, but that this result may be partially due to its high sampling intensity over time. Nevertheless, currently our results indicate that America’s newest national park may be a substantial exporter of free, native pollinators into economically-valuable agricultural lands as well as neighboring semi-wild lands. Only by comparing natural and disturbed areas over time to quantify the relative impacts of activities such as urbanization and agricultural intensification separate from more pervasive pressures like climate change, as is a goal of climate change vulnerability assessments [82], will we be able to determine the best multi-pronged approach to mitigating native bee declines. Our discovery that Pinnacles is the only area to have been extensively and repeatedly surveyed for bee biodiversity over multiple decades further underscores our call for increased repeated monitoring efforts to facilitate research on bee population decline and variability at its source.

## Acknowledgements

We are grateful to Therese Lamperty for dedication in the field, and to Harold Ikerd and Skyler Burrows for assistance in the lab. This work would not have been possible without generous guidance from USU co-P.I. Edward W. Evans and from Pinnacles wildlife biologist Paul G. Johnson. We thank Michael Orr, Skyler Burrows, Harold Ikerd, Karen Wright, Zachary Portman, Brian Rozick, and Ethan Frehner for help with bee identifications; Valerie Nuttman, Brent Johnson, and Denise Louie for support at Pinnacles; Amy Fesnock for her 2002 work on Pinnacles bees; Morgan Ernest, Paul Johnson, Eugene Schupp, Hao Ye, Erica Christensen, and Kenny Anderson for comments on drafts; Jereme Gaeta, Cody Griffin, and Audrey Wilson for assistance with figures; and our PlosONE editor for instructions that improved the manuscript.

## Supporting Information

**Table S1.** Pinnacles National Park long-term bee monitoring site details.

| **Site/Plot Number** | **Site/Plot Name** | **GPS lat/long** | **Elevation** | **Dimensions** | **Habitat Type** |
| --- | --- | --- | --- | --- | --- |
| 1 | McCabe Canyon Upper | 36.5081, -121.156 | 410m | 175 m x 57m | Alluvial |
| 2 | McCabe Canyon Lower | 36.503, -121.156 | 395m | 175m x 57m | Alluvial |
| 3 | Peaks View | 36.4802, -121.16 | 290m | 200m x 50m | Alluvial |
| 4 | South Wilderness | 36.4683, -121.156 | 280m | 250m x 40m | Live Oak Woodland |
| 5 | Needlegrass BOW | 36.5091, -121.12 | 385m | 200m x 50m | Blue Oak Woodland |
| 6 | Needlegrass LOW | 36.509, -121.129 | 365m | 200m x 50m | Live Oak Woodland |
| 7 | West Gate | 36.4747, -121.227 | 610m | 175m x 57m | Blue Oak Woodland |
| 8 | Double Gates | 36.4858, -121.232 | 535m | 200m x 50m | Grassland |
| 9 | W. North Wilderness | 36.4949, -121.211 | 430m | 200m x 50m | Grassland |
| 10 | High Peaks | 36.4907, -121.183 | 595m | 175m x 50m | Blue Oak Woodland |

**Fig. S1.**
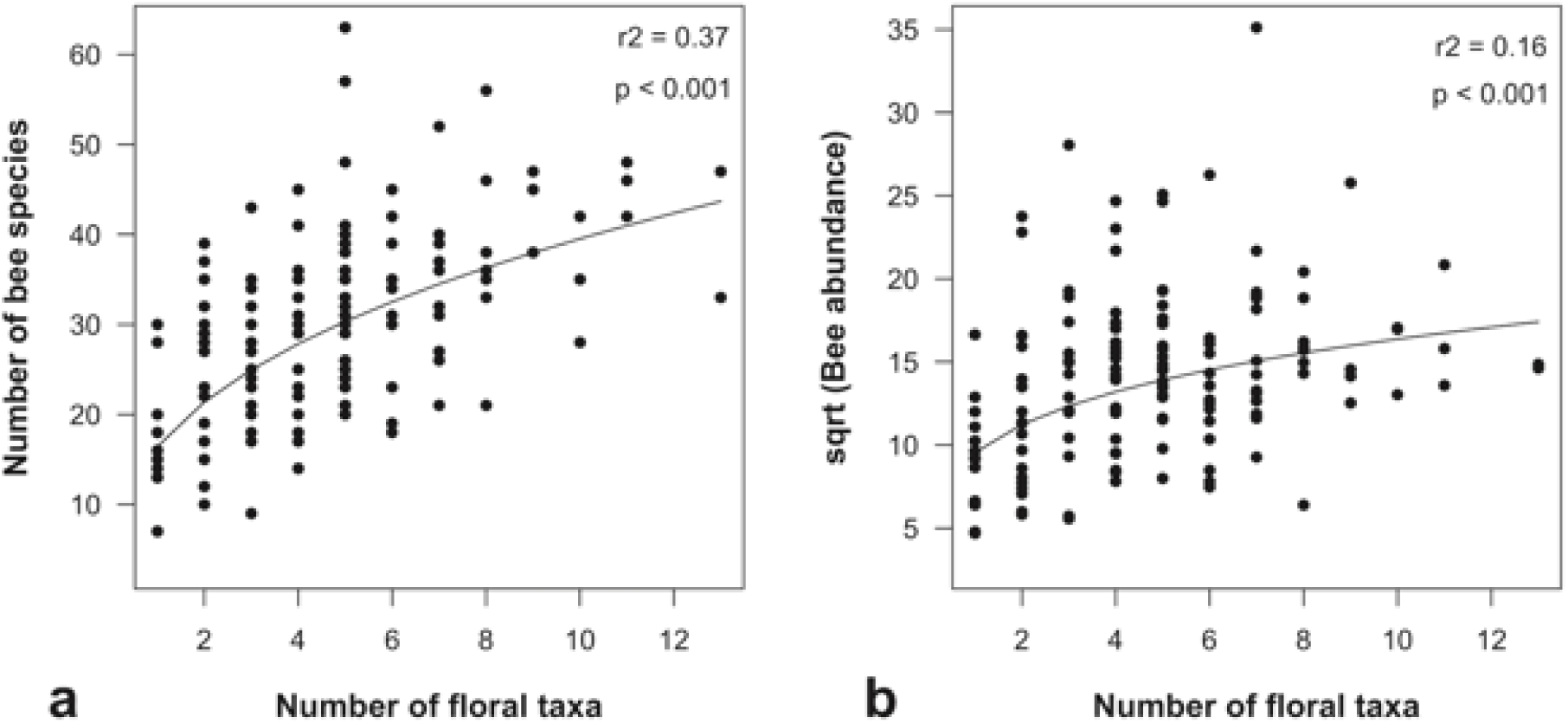
Relationship between floral richness (FR) and either (a) bee richness (BR) or (b) bee abundance (BA, square-root transformed) at the plot-sample level (N=150) within Pinnacles National Park (2011-12). Shown with power-law model (black line; (a) BR = exp(2.79 + 0.38*log(FR)); R^2^=0.37, p<0.01; (b) BA = exp(2.26 + 0.23*log(FR)); R^2^ = 0.16, p<0.01).

**Table S2.**
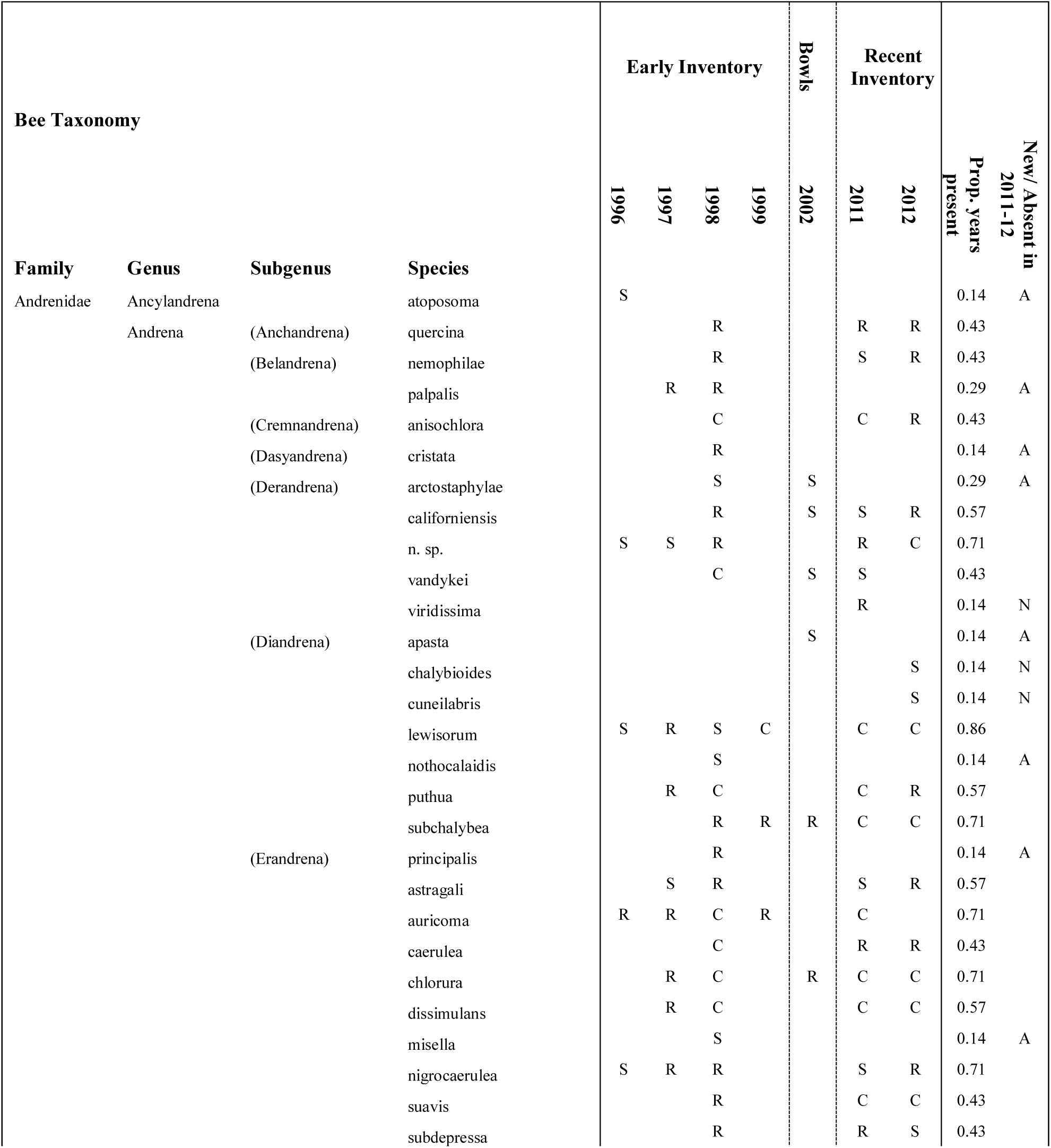
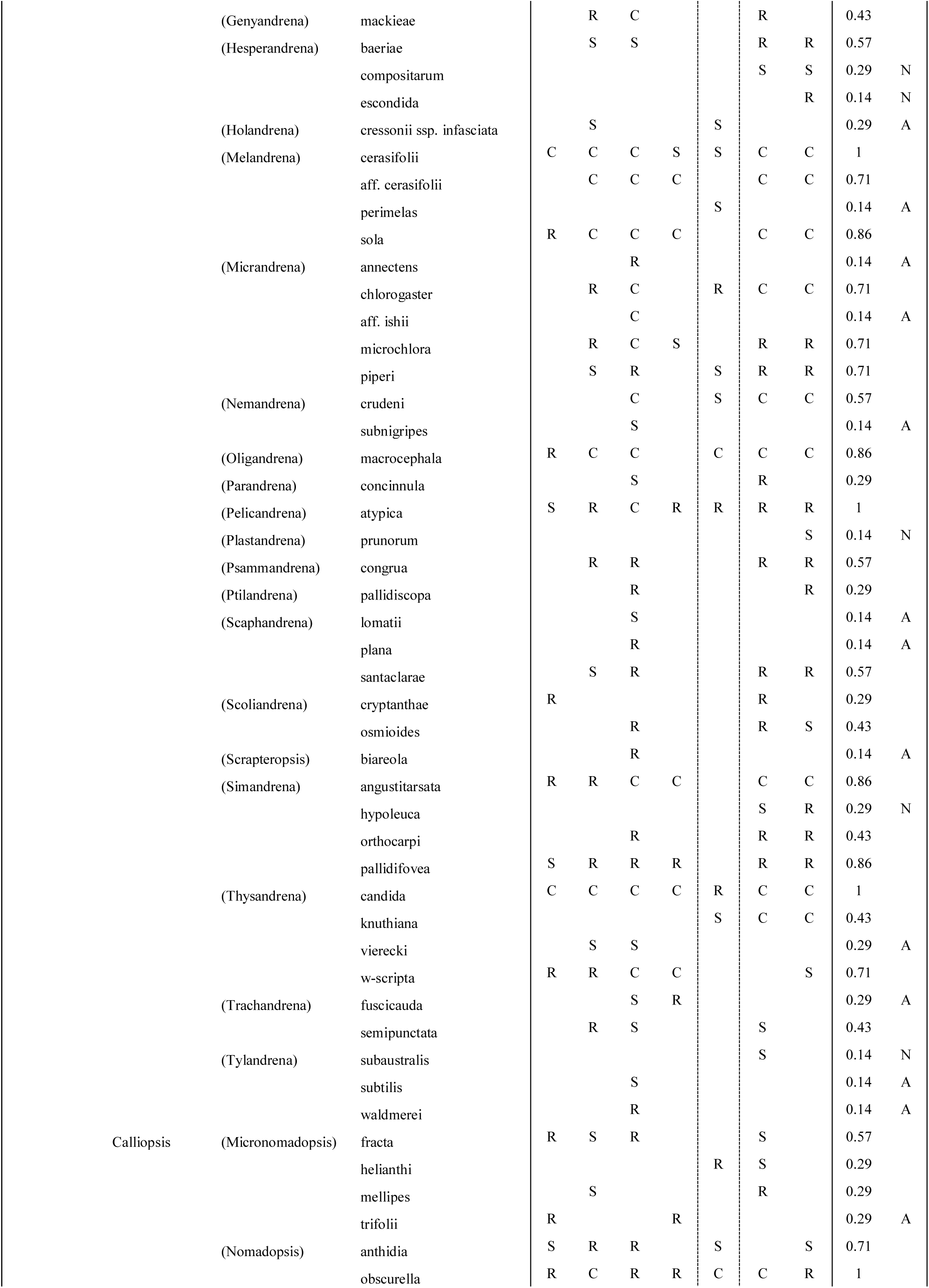
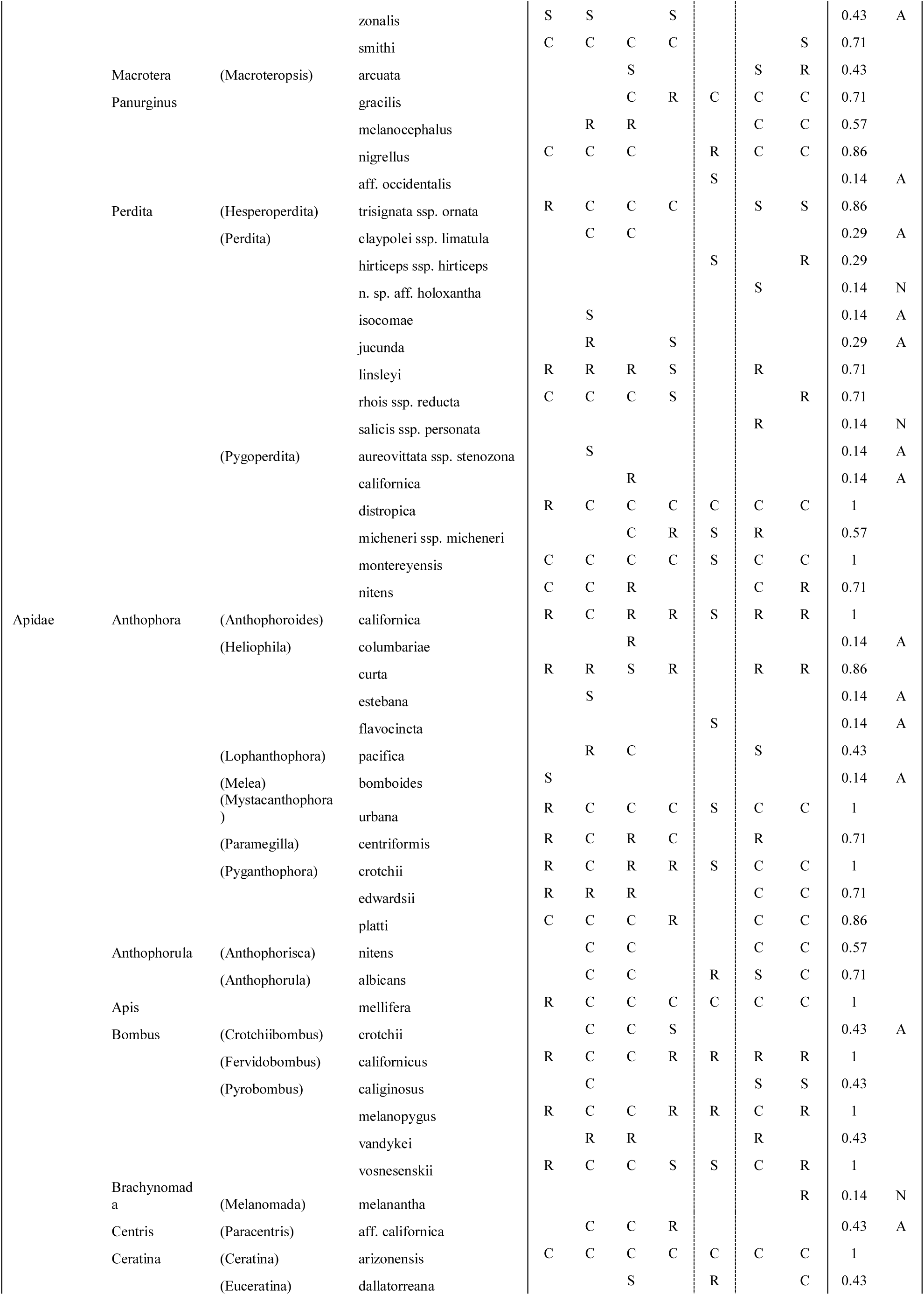
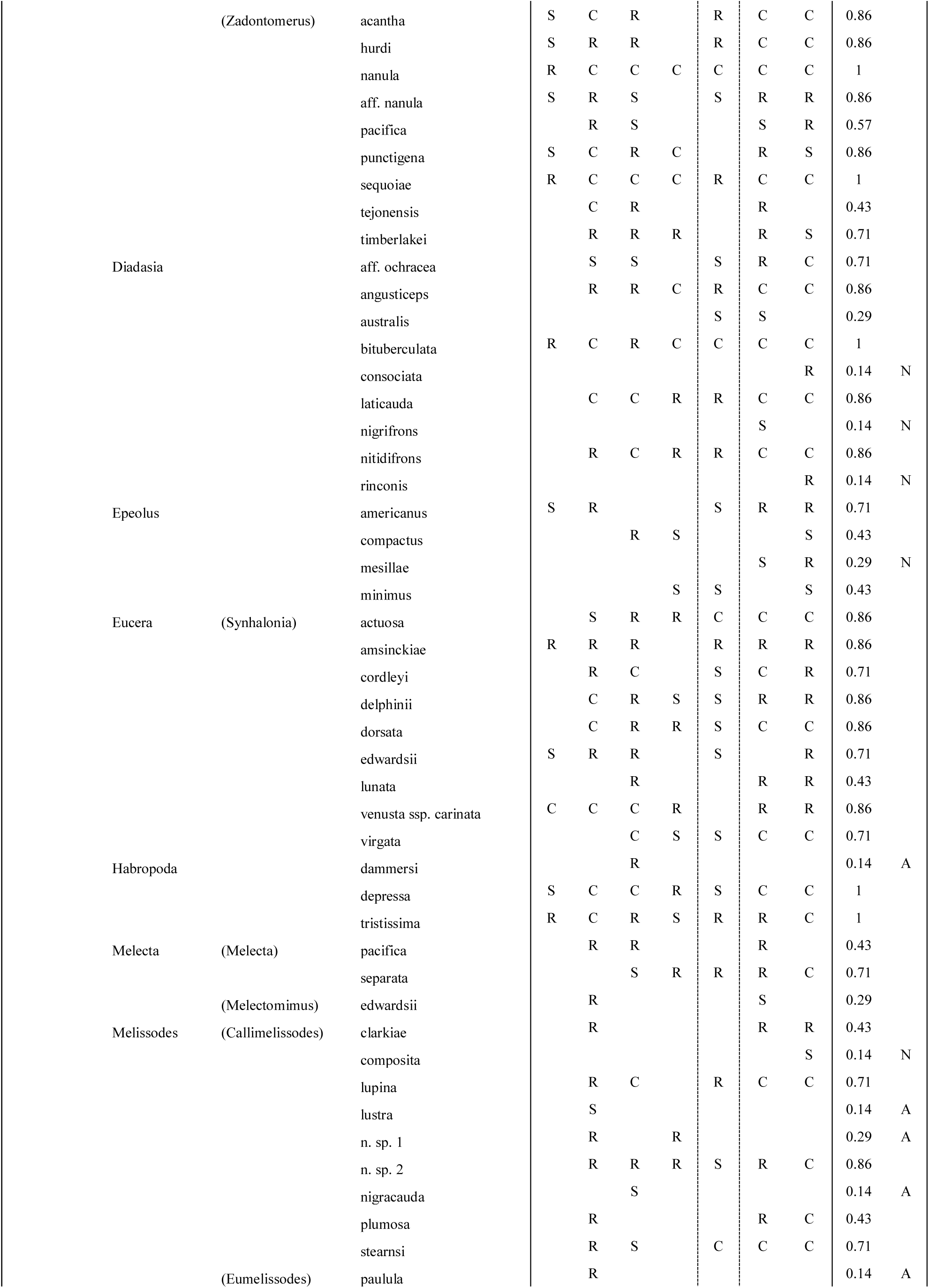
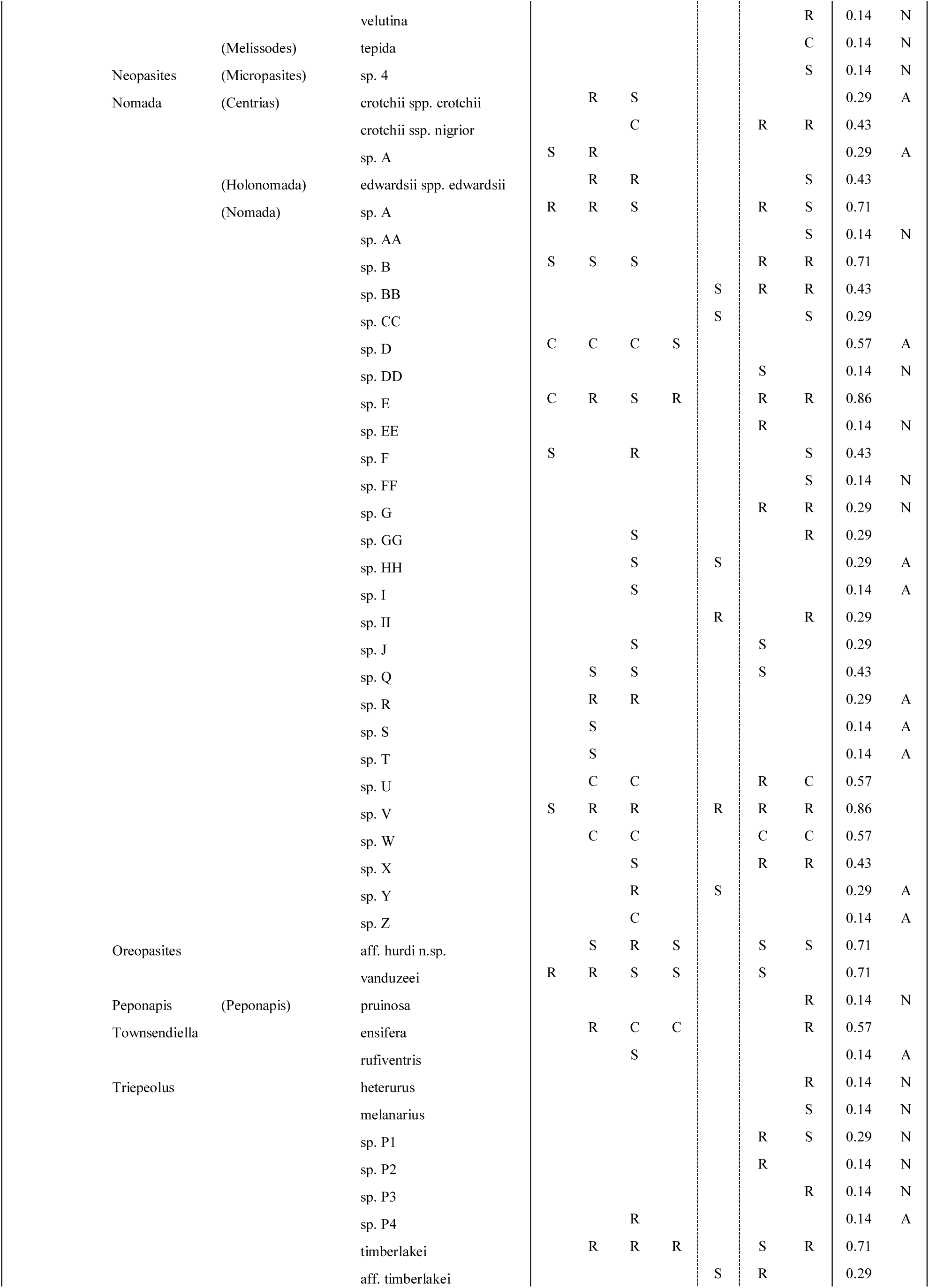
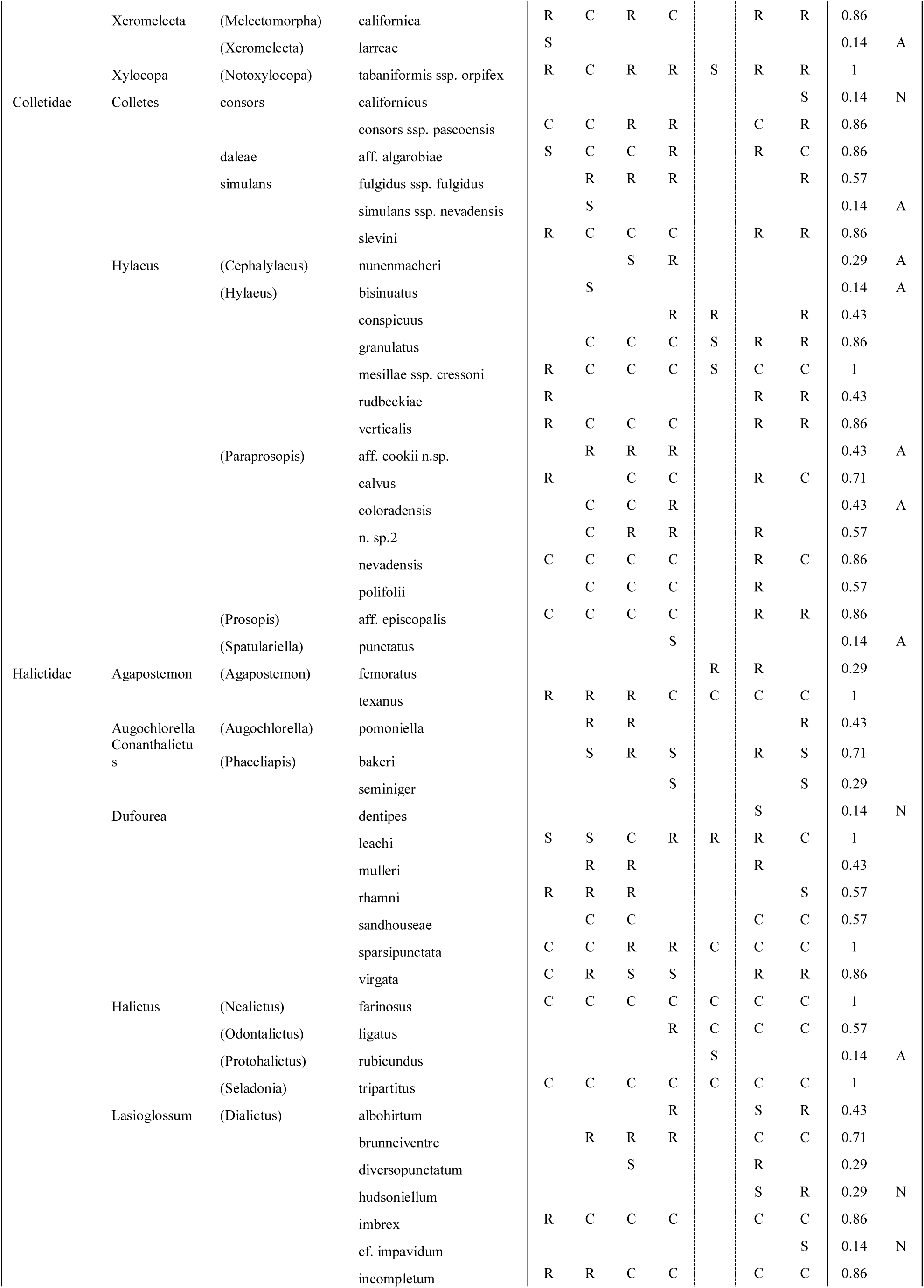
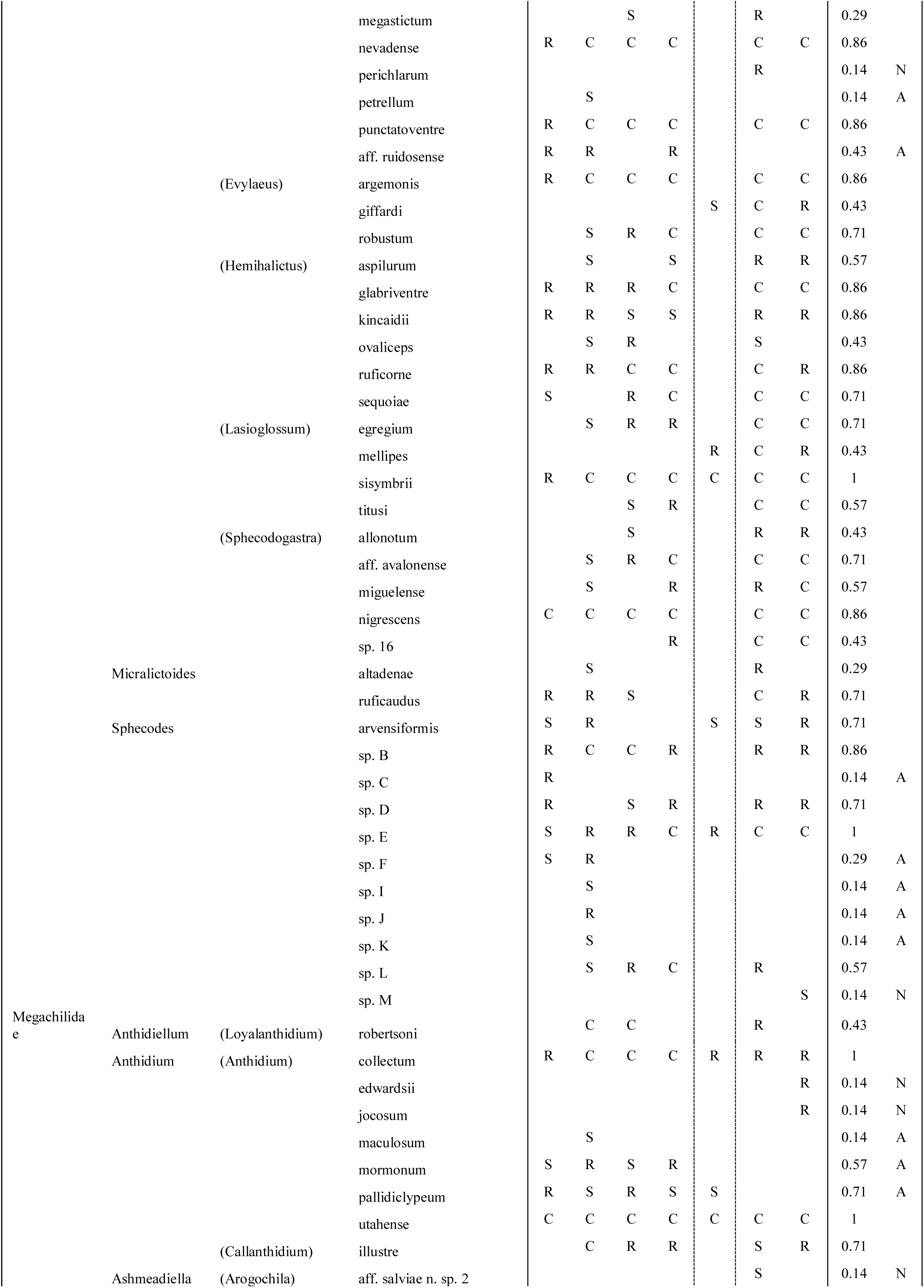
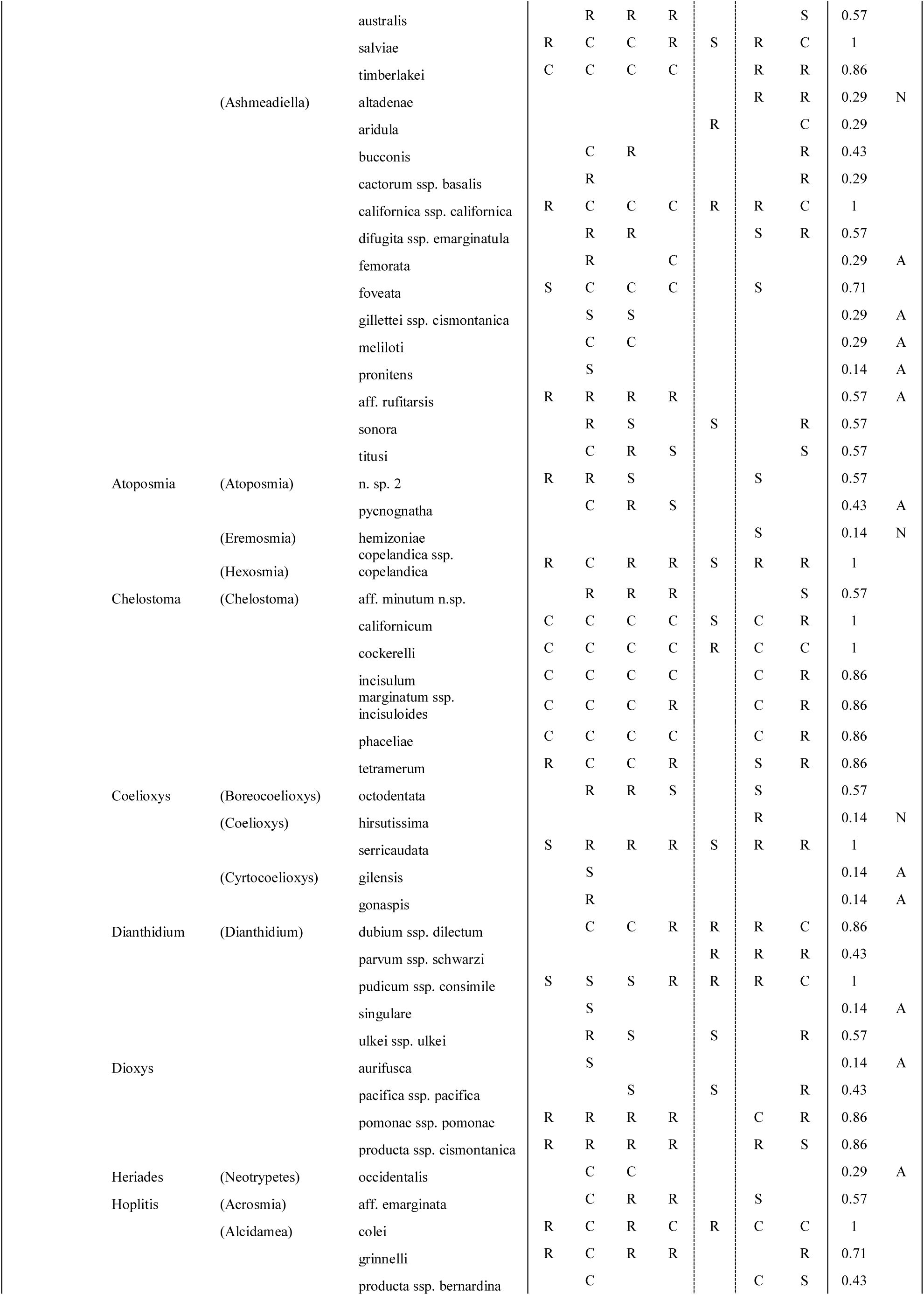
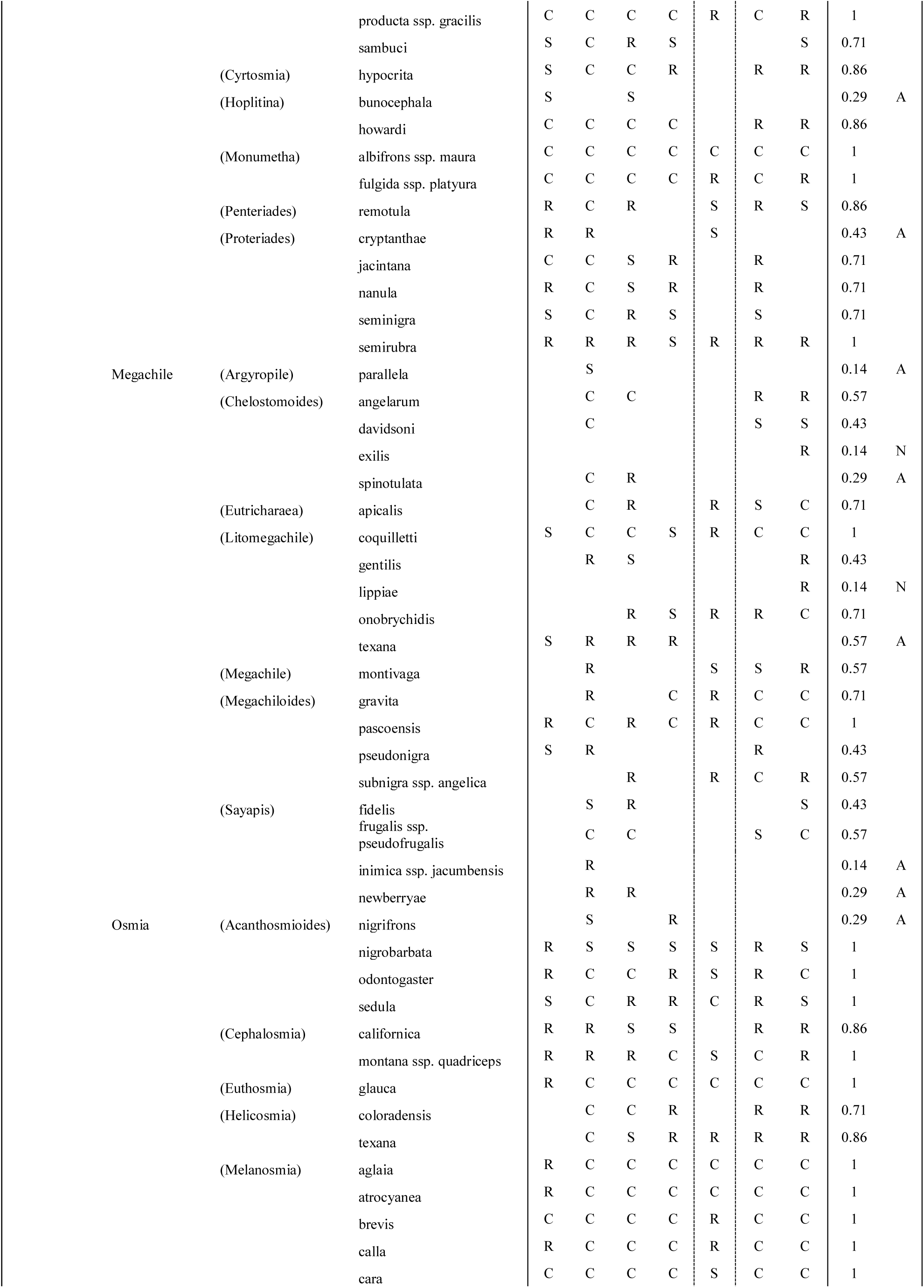
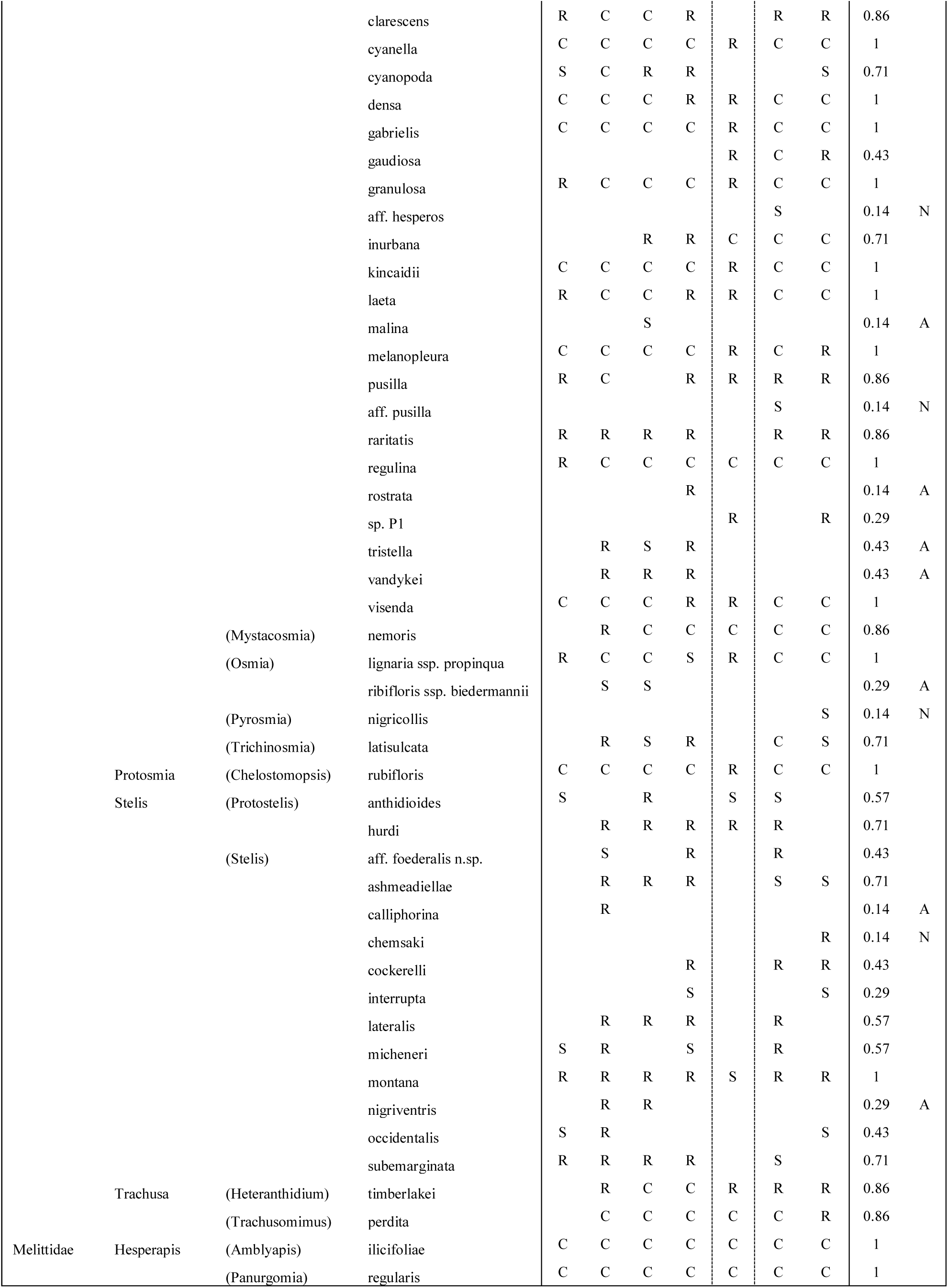
Full Pinnacles National Park bee species list, with relative abundance for each of seven collection years, proportion of years collected, and status as new (N) to or absent (A) from the current study. Species are marked “S” for Singleton if only one specimen was collected, “R” for Rare if N≤10, and “C” for Common if N>10. Dashed vertical line marks 2002 collection as separate from original 1996-9 inventory, but still prior to recent study (2011-12).

**Table S3.**
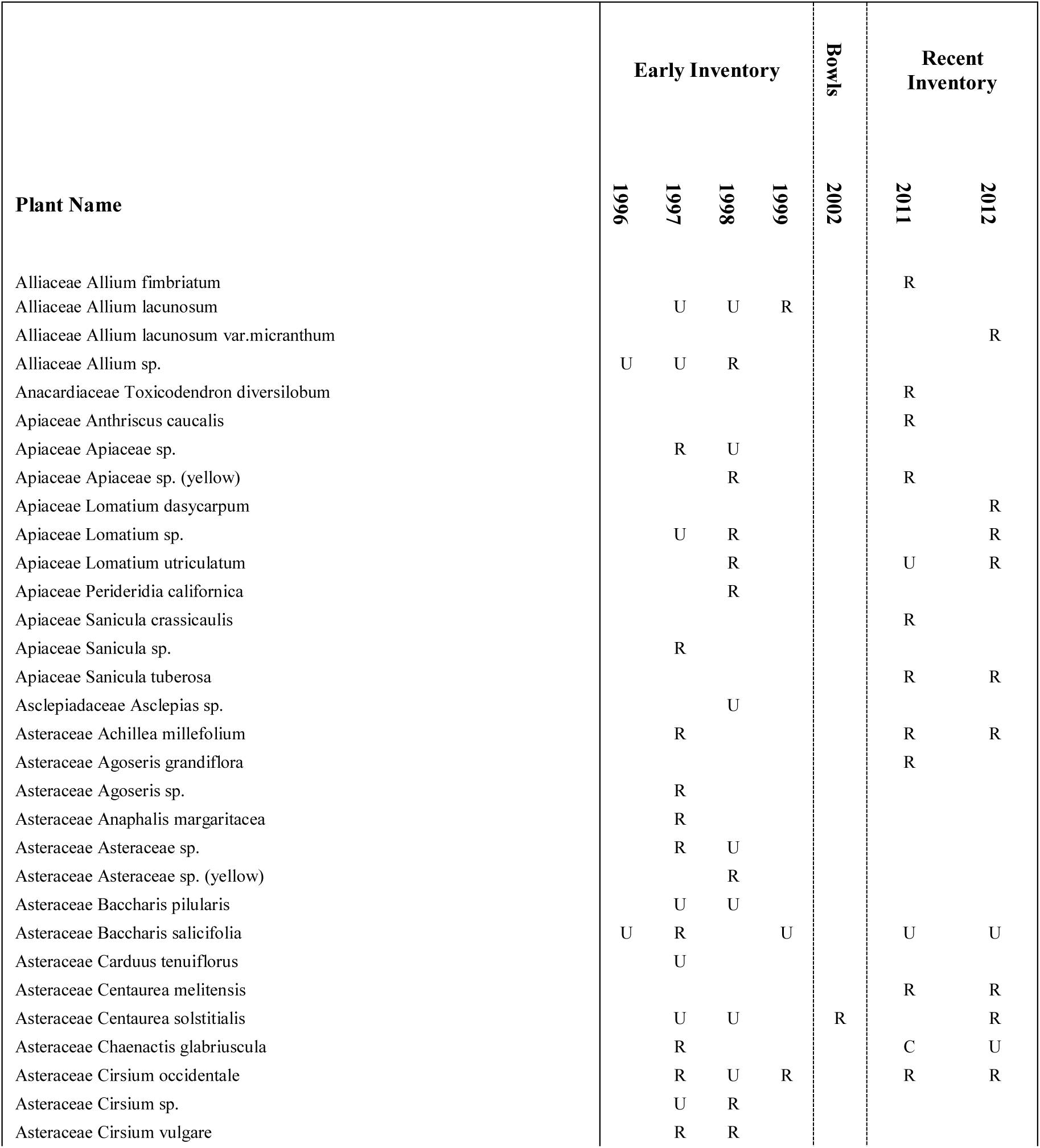
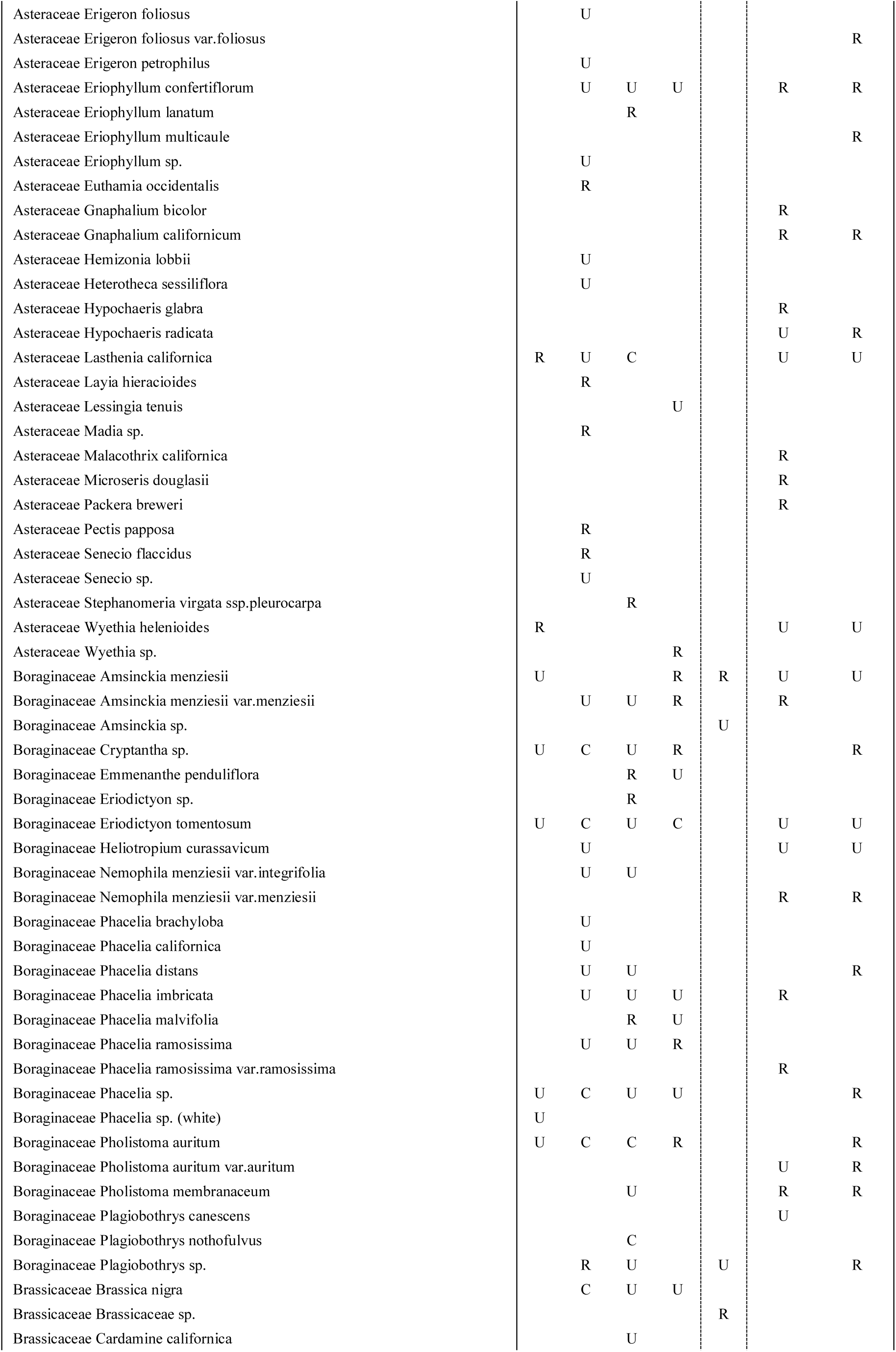
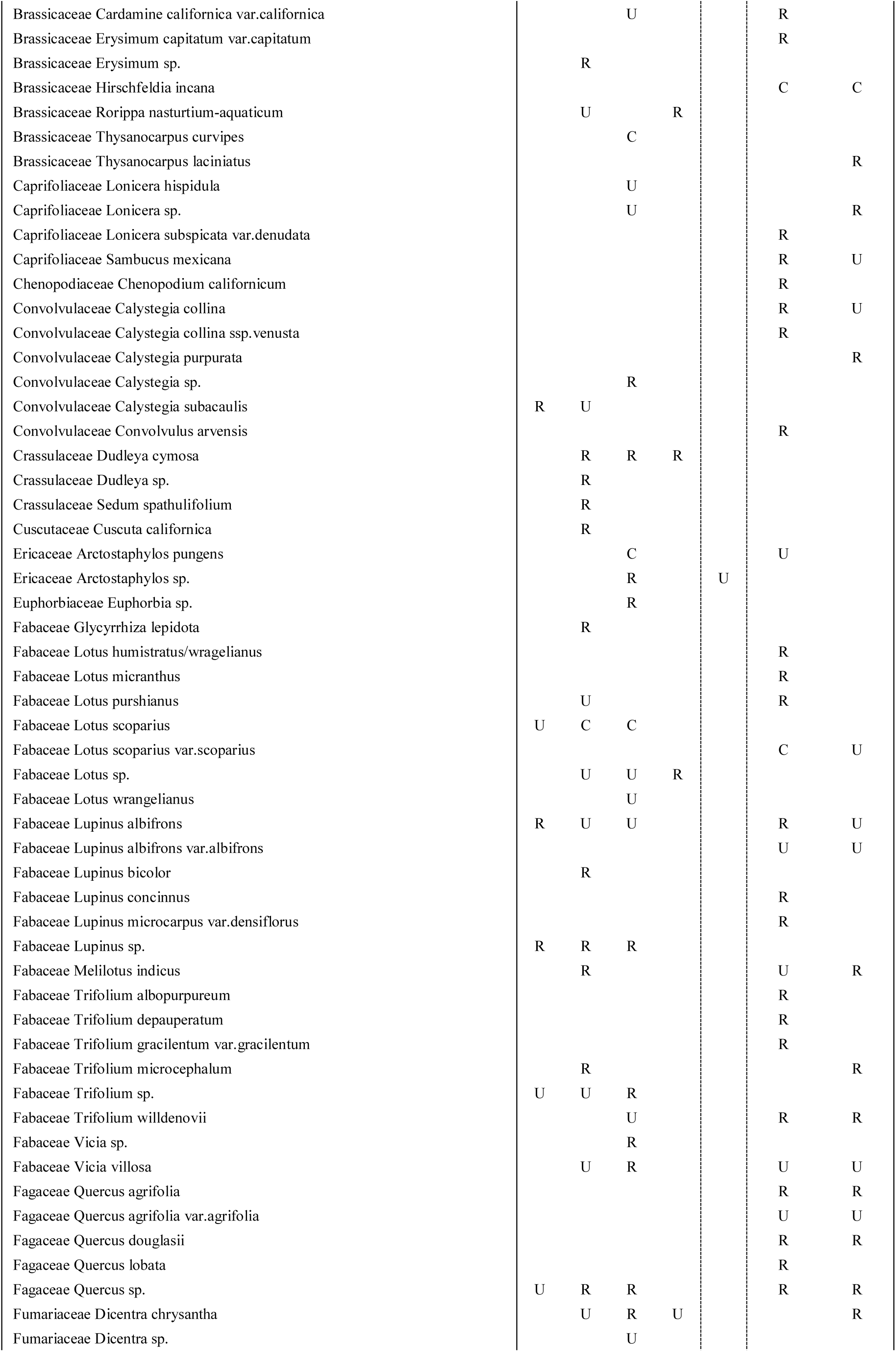
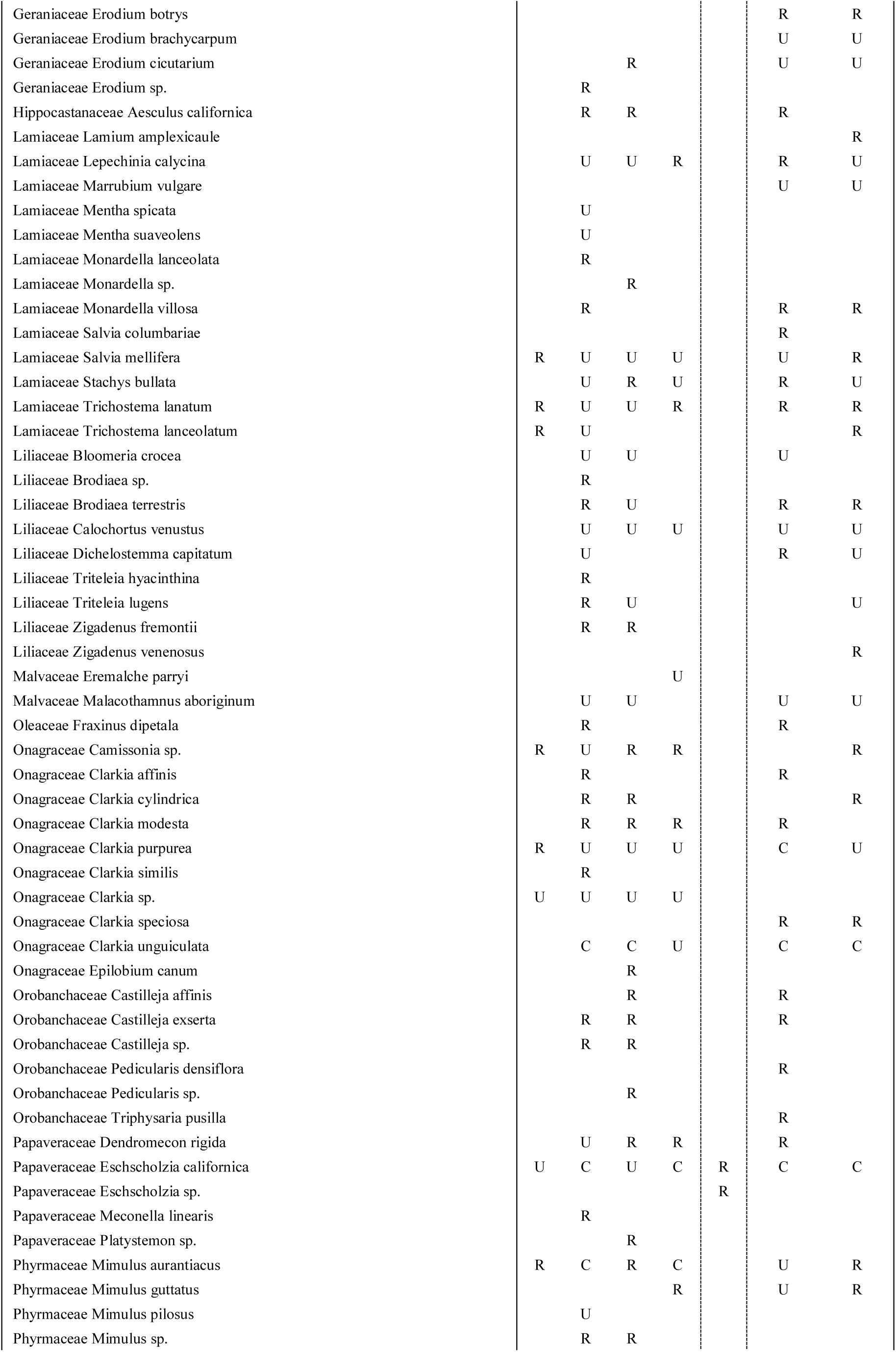
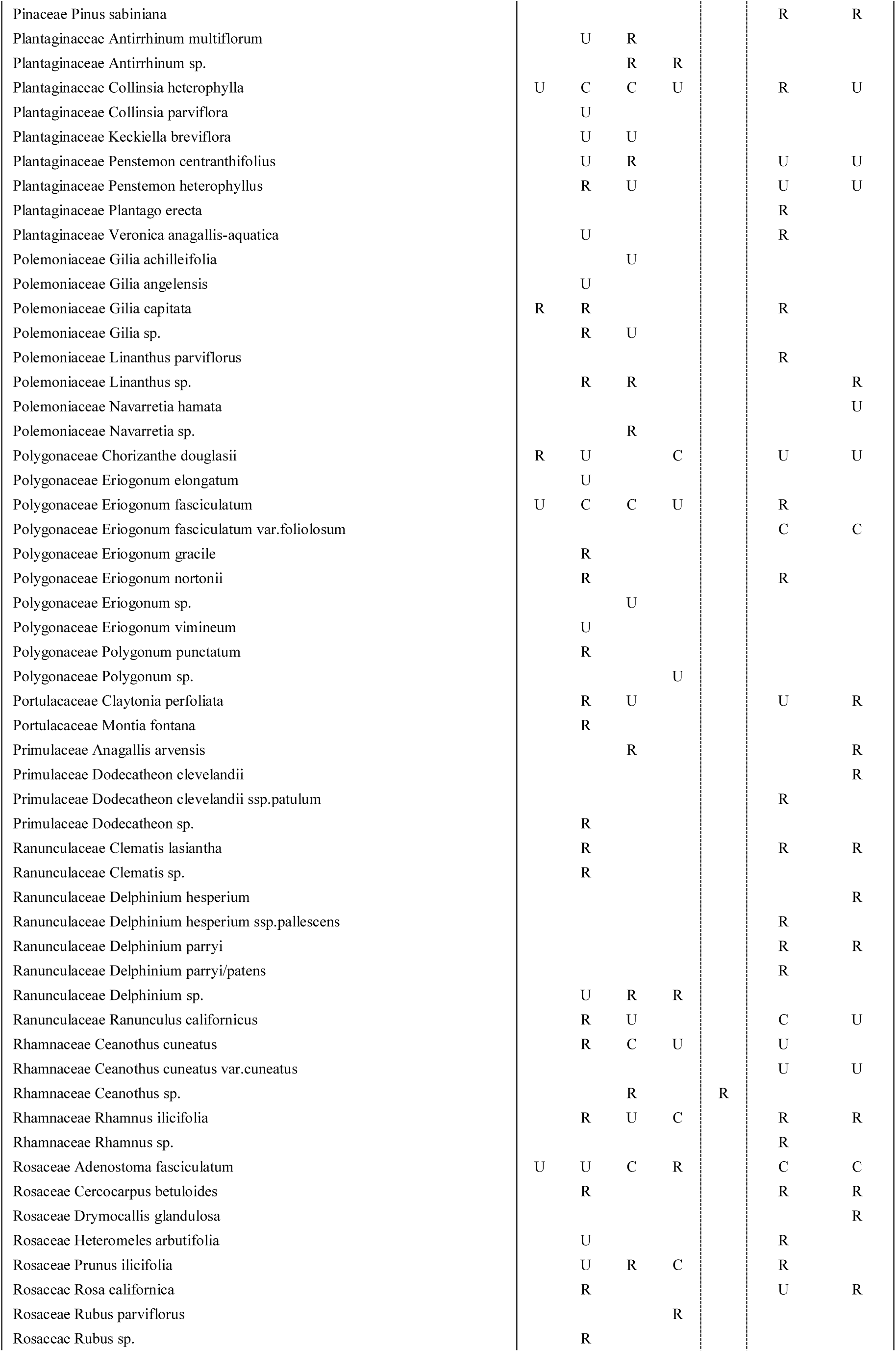
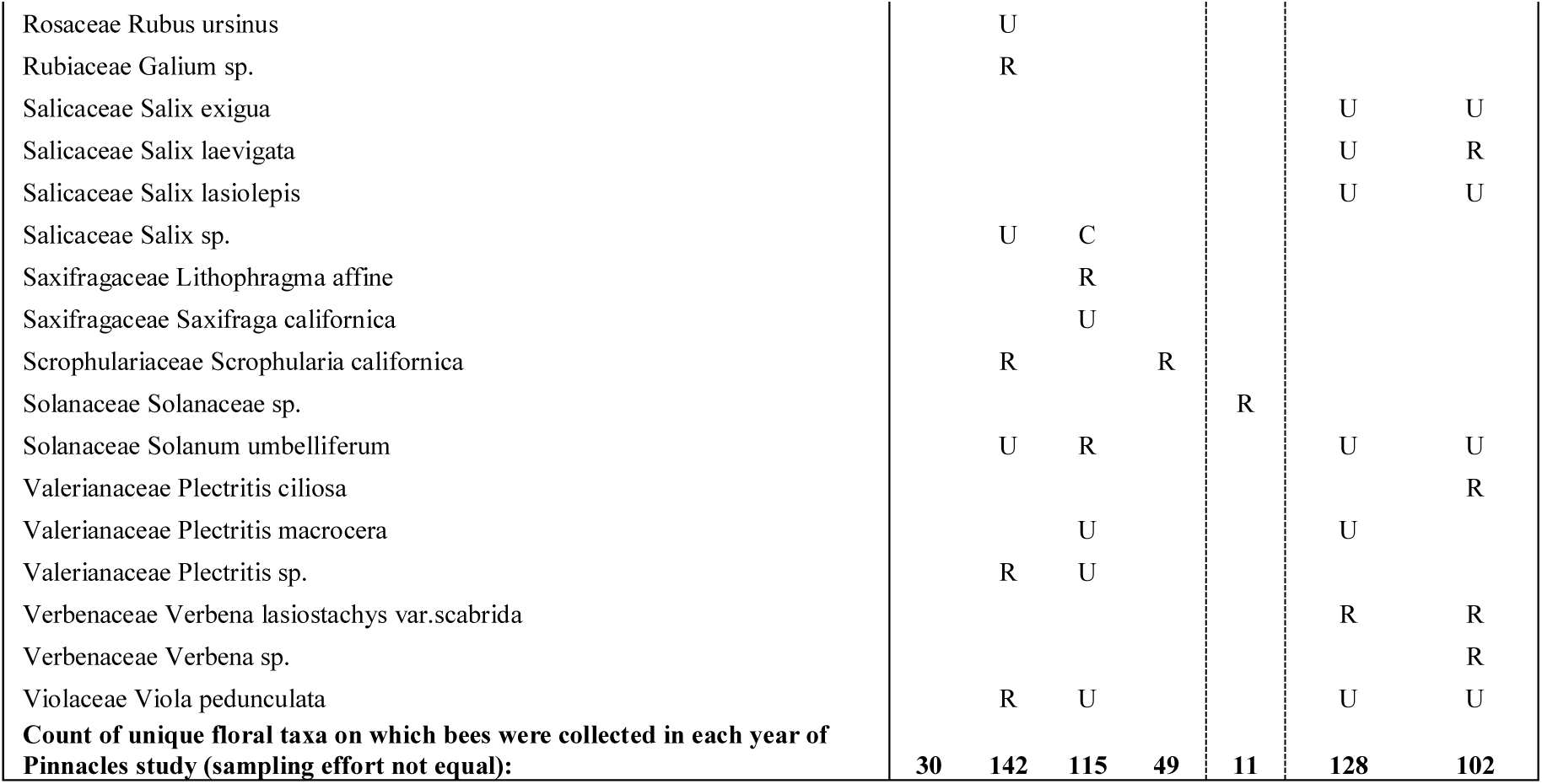
Floral taxa visited by bees at Pinnacles National Park (unique groups, identified to lowest possible level), and their relative popularity by year. Plants are marked with “R” for rare if bee visits were fewer than 10 in that year, with “U” for uncommon if bee visits ranged between 10-100, and “C” for common when over 100 bees were collected on that plant. The last row sums the plant taxa on which bees were collected per year. Dashed vertical line marks 2002 collection as separate from original 1996-9 study, and prior to the current study.

## References

1. Ollerton J, Winfree R, Tarrant S. How many flowering plants are pollinated by animals? Oikos. 2011;120: 321–326. doi:10.1111/j.1600-0706.2010.18644.x

2. Michener CD. The Bees of the World. Baltimore: Johns Hopkins University Press; 2007.

3. Morse R, Calderone NW. The value of honey bees as pollinators of U.S. Crops in 2000. Bee Cult. 2001;128. Available: http://agris.fao.org/agris-search/search.do?recordID=XE20122002449

4. Losey JE, Vaughan M. The Economic Value of Ecological Services Provided by Insects. BioScience. 2006;56: 311–323. doi:10.1641/0006-3568(2006)56[311:TEVOES]2.0.CO;2

5. Winfree R, Williams NM, Dushoff J, Kremen C. Native bees provide insurance against ongoing honey bee losses. Ecol Lett. 2007;10: 1105–1113. doi:10.1111/j.1461-0248.2007.01110.x

6. Garibaldi LA, Steffan-Dewenter I, Winfree R, Aizen MA, Bommarco R, Cunningham SA, et al. Wild Pollinators Enhance Fruit Set of Crops Regardless of Honey Bee Abundance. Science. 2013;339: 1608–1611. doi:10.1126/science.1230200

7. Greenleaf SS, Kremen C. Wild bees enhance honey bees’ pollination of hybrid sunflower. Proc Natl Acad Sci. 2006;103: 13890–13895. doi:10.1073/pnas.0600929103

8. Tepedino VJ. The importance of bees and other insect pollinators in maintaining floral species composition. Gt Basin Nat Mem. 1979; 139–150.

9. Potts SG, Vulliamy B, Dafni A, Ne’eman G, Willmer P. Linking Bees and Flowers: How Do Floral Communities Structure Pollinator Communities? Ecology. 2003;84: 2628–2642. doi:10.2307/3450108

10. Cane JH, Minckley RL, Kervin LJ, Roulston TH, Williams NM. Complex responses within a desert bee guild (Hymenoptera?: Apiformes) to urban habitat fragmentation. Ecol Appl. 2006;16: 632–644. doi:10.1890/1051-0761(2006)016[0632:CRWADB]2.0.CO;2

11. Kremen C, Williams NM, Bugg RL, Fay JP, Thorp RW. The area requirements of an ecosystem service: crop pollination by native bee communities in California. Ecol Lett. 2004;7: 1109–1119. doi:10.1111/j.1461-0248.2004.00662.x

12. Tscharntke T, Steffan-Dewenter I, Kruess A, Thies C. Characteristics of insect populations on habitat fragments: A mini review. Ecol Res. 2002;17: 229–239. doi:10.1046/j.1440-1703.2002.00482.x

13. Öckinger E, Smith HG. Semi-natural grasslands as population sources for pollinating insects in agricultural landscapes. J Appl Ecol. 2007;44: 50–59. doi:10.1111/j.1365-2664.2006.01250.x

14. Morandin LA, Kremen C. Hedgerow restoration promotes pollinator populations and exports native bees to adjacent fields. Ecol Appl. 2013;23: 829–839. doi:10.1890/12-1051.1

15. Chaplin-Kramer R, Dombeck E, Gerber J, Knuth KA, Mueller ND, Mueller M, et al. Global malnutrition overlaps with pollinator-dependent micronutrient production. Proc R Soc B. 2014;281: 20141799. doi:10.1098/rspb.2014.1799

16. Goulson D, Nicholls E, Botías C, Rotheray EL. Bee declines driven by combined stress from parasites, pesticides, and lack of flowers. Science. 2015;347: 1255957. doi:10.1126/science.1255957

17. Potts SG, Biesmeijer JC, Kremen C, Neumann P, Schweiger O, Kunin WE. Global pollinator declines: trends, impacts and drivers. Trends Ecol Evol. 2010;25: 345–353. doi:10.1016/j.tree.2010.01.007

18. Gonzalez VH, Griswold T, Engel MS. Obtaining a better taxonomic understanding of native bees: where do we start? Syst Entomol. 2013;38: 645–653. doi:10.1111/syen.12029

19. Linsley EG. The Ecology of Solitary Bees. Hilgardia. 1958;27: 543–599.

20. Michener CD. Biogeography of the bees. Ann Mo Bot Gard. 1979;66: 277–347.

21. Cardoso P, Erwin TL, Borges PAV, New TR. The seven impediments in invertebrate conservation and how to overcome them. Biol Conserv. 2011;144: 2647–2655. doi:10.1016/j.biocon.2011.07.024

22. Fischer AG. Latitudinal Variations in Organic Diversity. Evolution. 1960;14: 64–81. doi:10.2307/2405923

23. Minckley R. Faunal composition and species richness differences of bees (Hymenoptera: Apiformes) from two north American regions. Apidologie. 2008;39: 176–188. doi:10.1051/apido:2007062

24. Minckley RL, Roulston TH, Williams NM. Resource assurance predicts specialist and generalist bee activity in drought. Proc R Soc B Biol Sci. 2013;280. doi:10.1098/rspb.2012.2703

25. Williams N, Minckley R, Silveira F. Variation in native bee faunas and its implications for detecting community changes. Conserv Ecol. 2001;5: 1–24.

26. Bascompte J, Jordano P, Olesen J. Asymmetric coevolutionary networks facilitate biodiversity maintenance. Science. 2006;312: 431–433. doi:10.1126/science.1123412

27. Bascompte J, Jordano P, Melián CJ, Olesen JM. The nested assembly of plant–animal mutualistic networks. Proc Natl Acad Sci. 2003;100: 9383–9387. doi:10.1073/pnas.1633576100

28. Kremen C. Managing ecosystem services: what do we need to know about their ecology? Ecol Lett. 2005;8: 468–479. doi:10.1111/j.1461-0248.2005.00751.x

29. Larsen TH, Williams NM, Kremen C. Extinction order and altered community structure rapidly disrupt ecosystem functioning. Ecol Lett. 2005;8: 538–547. doi:10.1111/j.1461-0248.2005.00749.x

30. Williams NM, Crone EE, Roulston TH, Minckley RL, Packer L, Potts SG. Ecological and life-history traits predict bee species responses to environmental disturbances. Biol Conserv. 2010;143: 2280–2291. doi:10.1016/j.biocon.2010.03.024

31. Aizen MA, Sabatino M, Tylianakis JM. Specialization and Rarity Predict Nonrandom Loss of Interactions from Mutualist Networks. Science. 2012;335: 1486–1489. doi:10.1126/science.1215320

32. Bartomeus I, Ascher JS, Gibbs J, Danforth BN, Wagner DL, Hedtke SM, et al. Historical changes in northeastern US bee pollinators related to shared ecological traits. Proc Natl Acad Sci. 2013;110: 4656–4660. doi:10.1073/pnas.1218503110

33. Bommarco R, Biesmeijer JC, Meyer B, Potts SG, Pöyry J, Roberts SPM, et al. Dispersal capacity and diet breadth modify the response of wild bees to habitat loss. Proc R Soc Lond B Biol Sci. 2010; rspb20092221. doi:10.1098/rspb.2009.2221

34. Memmott J, Craze PG, Waser NM, Price MV. Global warming and the disruption of plant–pollinator interactions. Ecol Lett. 2007;10: 710–717. doi:10.1111/j.1461-0248.2007.01061.x

35. Forrest JRK, Thomson JD. An examination of synchrony between insect emergence and flowering in Rocky Mountain meadows. Ecol Monogr. 2011;81: 469–491. doi:10.1890/10-1885.1

36. Vilela AA, Del Claro VTS, Torezan-Silingardi HM, Del-Claro K. Climate changes affecting biotic interactions, phenology, and reproductive success in a savanna community over a 10-year period. Arthropod-Plant Interact. 2018;12: 215–227. doi:10.1007/s11829-017-9572-y

37. Meiners JM, Griswold TL, Harris DJ, Ernest SKM. Bees without Flowers: Before Peak Bloom, Diverse Native Bees Find Insect-Produced Honeydew Sugars. Am Nat. 2017;190: 281–291. doi:10.1086/692437

38. Romero GQ, Gonçalves-Souza T, Kratina P, Marino NAC, Petry WK, Sobral-Souza T, et al. Global predation pressure redistribution under future climate change. Nat Clim Change. 2018;8: 1087. doi:10.1038/s41558-018-0347-y

39. Cornelissen T. Climate change and its effects on terrestrial insects and herbivory patterns. Neotrop Entomol. 2011;40: 155–163. doi:10.1590/S1519-566X2011000200001

40. Griswold TL, Andres M, Andrus R, Garvin G, Keen K, Kervin L, et al. A survey of the rare bees of Clark County, Nevada. Final Rep Nat Conserv Las Vegas NV. 1999; Available: http://works.bepress.com/terry_griswold/63

41. Marlin JC, LaBerge WE. The Native Bee Fauna of Carlinville, Illinois, Revisited After 75 Years:a Case for Persistence. Ecol Soc. 2007; Available: http://agris.fao.org/agris-search/search.do?recordID=XE20122002329

42. Messinger O, Griswold TL. A Pinnacle of bees. Fremontia. 2003;30: 32–40.

43. Messinger O. A survey of the bees of Grand Staircase-Escalante National Monument, Southern Utah: Incidence, Abundance, and Community dynamics. Masters of Science, Utah State University. 2006.

44. Roubik DW. Ups and downs in pollinator populations: When is there a decline? 2001; Available: http://dlc.dlib.indiana.edu/dlc/handle/10535/3364

45. Wilson JS, Messinger OJ, Griswold T. Variation between bee communities on a sand dune complex in the Great Basin Desert, North America: Implications for sand dune conservation. J Arid Environ. 2009;73: 666–671.

46. Drons DJ. An Inventory of Native Bees (Hymenoptera: Apiformes) in the Black Hills of South Dakota and Wyoming [Internet]. South Dakota State University. 2012. Available: http://gfp.sd.gov/images/WebMaps/Viewer/WAP/Website/SWGSummaries/Drons%202012_an%20inventory%20of%20native%20Black%20Hills%20bees%20acknowledge.pdf

47. Giles V, Ascher JS. A survey of the bees of the Black Rock Forest preserve, New York (Hymenoptera: Apoidea). J Hymenopt Res. 2006;15: 208–231.

48. Moldenke AR. California pollination ecology and vegetation types. Phytologia. 1976;34: 305–361.

49. Moldenke AR. Evolutionary history and diversity of the bee faunas of Chile and Pacific North America. Wasmann J Biol. 1976;34: 147–178.

50. Meiners JM, Griswold TL, Evans EW. Native Bees of Pinnacles National Park: Diversity Inventory and Plot Sampling Final Report and Sampling Manual. National Park Service; Utah State University; 2015 Sep p. 76.

51. Meiners JM. Biodiversity, Community Dynamics, and Novel Foraging Behaviors of a Rich Native Bee Fauna across Habitats at Pinnacles National Park, California. Masters of Science, Utah State University. 2016.

52. LeBuhn G, Griswold T, Minckley R, Droege S, Roulston T, Cane J, et al. A standardized method for monitoring bee populations–the bee inventory (BI) plot. 2003; Available: http://cybercemetery.unt.edu/archive/nbii/20120111121317/http://online.sfsu.edu/∼beeplot/pdfs/Bee%20Plot%202003.pdf

53. Baldwin BG, Goldman DH. The Jepson Manual: Vascular Plants of California. University of California Press; 2012.

54. R Core Team. R: A language and environment for statistical computing. https://www.R-project.org. Vienna, Austria; 2015.

55. Oksanen J, Blanchet FG, Friendly M, Kindt R, Legendre P, McGlinn D, et al. vegan: Community Ecology Package [Internet]. 2018. Available: https://CRAN.R-project.org/package=vegan

56. Arrhenius O. Species and Area. J Ecol. 1921;9: 95–99. doi:10.2307/2255763

57. Connor EF, McCoy ED. The Statistics and Biology of the Species-Area Relationship. Am Nat. 1979;113: 791–833. doi:10.1086/283438

58. Gotelli NJ, Colwell RK. Quantifying biodiversity: procedures and pitfalls in the measurement and comparison of species richness. Ecol Lett. 2001;4: 379–391. doi:10.1046/j.1461-0248.2001.00230.x

59. Bohart G, Knowlton G. The Bees of Curlew Valley (Utah and Idaho). Proc Utah Acad Sci Arts Lett. 1973; Available: http://digitalcommons.usu.edu/piru_pubs/790

60. Griswold T, Parker F, Tepedino V. The bees of the San Rafael Desert: Implications for the bee fauna of the Grand Staircase-Escalante National Monument. In: Hill LM, editor. Learning from the land: Grand Staircase Escalante National Monument Science Symposium Proceedings. 1998. pp. 175–186.

61. Brown JW, Bahr SM. The Insect (Insecta) Fauna of Plummers Island, Maryland: Brief Collecting History and Status of the Inventory. Bull Biol Soc Wash. 2008;15: 54–64. doi:10.2988/0097-0298(2008)15[54:TIIFOP]2.0.CO;2

62. Kuhlman M, Burrows S. Checklist of bees (Apoidea) from a private conservation property in west-central Montana. Biodivers Data J.

63. Grundel R, Jean RP, Frohnapple KJ, Gibbs J, Glowacki GA, Pavlovic NB. A Survey of Bees (Hymenoptera: Apoidea) of the Indiana Dunes and Northwest Indiana, USA. J Kans Entomol Soc. 2011;84: 105–138. doi:10.2317/JKES101027.1

64. Tepedino VJ, Stanton NL. Diversity and Competition in Bee-Plant Communities on Short-Grass Prairie. Oikos. 1981;36: 35–44. doi:10.2307/3544376

65. Rhoades PR, Griswold T, Ikerd H, Waits L, Bosque-Pérez N, Eigenbrode S. The native bee fauna of the Palouse Prairie (Hymenoptera: Apoidea). J Melittology. 2017;0: 1–20. doi:10.17161/jom.v0i66.5703

66. Rust R, Menke A, Miller D. A biogeographic comparison of the bees, sphecid wasps, and mealybugs of the California Channel Ilsnads (Hymenoptera, Homoptera). Entomology of the California Channel Islands: proceedings of the first symposium. Santa Barbara, CA: Santa Barbara Museum of Natural History; 1985. pp. 22–59.

67. Wilson JS, Wilson LE, Loftis LD, Griswold T. The Montane Bee Fauna of North Central Washington, USA, with Floral Associations. West North Am Nat. 2010;70: 198–207. doi:10.3398/064.070.0206

68. Smith BA, Brown RL, Laberge W, Griswold T. A Faunistic Survey of Bees (Hymenoptera: Apoidea) in the Black Belt Prairie of Mississippi. J Kans Entomol Soc. 2012;85: 32–47. doi:10.2317/JKES111025.1

69. Deyrup M, Edirisinghe J, Norden B. The diversity and floral hosts of bees at the Archbold Biological Station, Florida (Hymenoptera: Apoidea). Available: http://publikationen.ub.uni-frankfurt.de/frontdoor/index/index/year/2013/docId/23868

70. Michener CD. Bees of a Limited Area in Southern Mississippi (Hymenoptera; Apoidea). Am Midl Nat. 1947;38: 443–455. doi:10.2307/2421575

71. Holt RD, Lawton JH, Polis GA, Martinez ND. Trophic Rank and the Species–Area Relationship. Ecology. 1999;80: 1495–1504. doi:10.1890/0012-9658(1999)080[1495:TRATSA]2.0.CO;2

72. Rosenzweig ML, L RM. Species Diversity in Space and Time. Cambridge University Press; 1995.

73. Frankie GW, Thorp RW, Hernandez J, Rizzardi M, Ertter B, Pawelek JC, et al. Native bees are a rich natural resource in urban California gardens. Calif Agric. 2009;63: 113–120. doi:10.3733/ca.v063n03p113

74. USDA - National Agricultural Statistics Service - Honey Bee Surveys and Reports [Internet]. [cited 20 Nov 2016]. Available: https://www.nass.usda.gov/Surveys/Guide_to_NASS_Surveys/Bee_and_Honey/

75. Kremen C, Ricketts T. Global Perspectives on Pollination Disruptions. Conserv Biol. 2000;14: 1226–1228.

76. Wolf AT, Ascher JS. Bees of Wisconsin (Hymenoptera: Apoidea: Anthophila). Gt Lakes Entomol. 2009;41.1: 129–168.

77. Donovan BJ. INTERACTIONS BETWEEN NATIVE AND INTRODUCED BEES IN NEW ZEALAND. N Z J Ecol. 1980;3: 104–116.

78. Petanidou T, Kallimanis AS, Tzanopoulos J, Sgardelis SP, Pantis JD. Long-term observation of a pollination network: fluctuation in species and interactions, relative invariance of network structure and implications for estimates of specialization. Ecol Lett. 2008;11: 564–575. doi:10.1111/j.1461-0248.2008.01170.x

79. Báldi A. Habitat heterogeneity overrides the species–area relationship. J Biogeogr. 2008;35: 675–681. doi:10.1111/j.1365-2699.2007.01825.x

80. Tews J, Brose U, Grimm V, Tielbörger K, Wichmann MC, Schwager M, et al. Animal species diversity driven by habitat heterogeneity/diversity: the importance of keystone structures: Animal species diversity driven by habitat heterogeneity. J Biogeogr. 2004;31: 79–92. doi:10.1046/j.0305-0270.2003.00994.x

81. Klausmeyer KR, Shaw MR. Climate Change, Habitat Loss, Protected Areas and the Climate Adaptation Potential of Species in Mediterranean Ecosystems Worldwide. PLoS ONE. 2009;4: e6392. doi:10.1371/journal.pone.0006392

82. Foden WB, Young BE, Akçakaya HR, Garcia RA, Hoffmann AA, Stein BA, et al. Climate change vulnerability assessment of species. Wiley Interdiscip Rev Clim Change. 0: e551. doi:10.1002/wcc.551

83. Matthews VI. Correlation of Pinnacles and Neenach Volcanic Formations and Their Bearing on San Andreas Fault Problem. AAPG Bull. 1976;60: 2128–2141.

84. Tucker S, Knudsen K, Robertson J. Additional lichen collections from Pinnacles National Monument, San Benito County, California. Bull Calif Lichen Soc. 2006;13: 8–11.

85. NPS. Nature - Pinnacles National Park (U.S. National Park Service) [Internet]. 2015 [cited 15 Nov 2015]. Available: http://www.nps.gov/pinn/learn/nature/index.htm

86. Debano SJ. Effects of livestock grazing on aboveground insect communities in semi-arid grasslands of southeastern Arizona. Biodivers Conserv. 2006;15: 2547. doi:10.1007/s10531-005-2786-9

87. Winfree R, Aguilar R, Vázquez DP, LeBuhn G, Aizen MA. A meta-analysis of bees’ responses to anthropogenic disturbance. Ecology. 2009;90: 2068–2076. doi:10.1890/08-1245.1

88. Iles DT, Williams NM, Crone EE. Source-sink dynamics of bumblebees in rapidly changing landscapes. J Appl Ecol. 2018;55: 2802–2811. doi:10.1111/1365-2664.13175

89. Griffin SR, Bruninga-Socolar B, Kerr MA, Gibbs J, Winfree R. Wild bee community change over a 26-year chronosequence of restored tallgrass prairie. Restor Ecol. 2017; doi:10.1111/rec.12481

90. Albrecht M, Riesen M, Schmid B. Plant-pollinator network assembly along the chronosequence of a glacier foreland. Oikos. 2010;119: 1610–1624. doi:10.1111/j.1600-0706.2010.18376.x

